# Correlative light and electron microscopy for human brain and other biological models

**DOI:** 10.1101/2024.06.11.598271

**Authors:** Notash Shafiei, Daniel Stӓhli, Domenic Burger, Marta Di Fabrizio, Lukas van den Heuvel, Jean Daraspe, Carolin Böing, Sarah H Shahmoradian, Wilma DJ van de Berg, Christel Genoud, Henning Stahlberg, Amanda J Lewis

**Affiliations:** Laboratory of Biological Electron Microscopy, Institute of Physics, School of Basic Sciences, Ecole Polytechnique Fédérale de Lausanne, 1015 Lausanne, Switzerland; Department of Fundamental Microbiology, Faculty of Biology and Medicine, University of Lausanne, 1015 Lausanne, Switzerland; Electron Microscopy Facility, Biophore, University of Lausanne, 1015 Lausanne, Switzerland; C-CINA, Biozentrum, University of Basel, 4058 Basel, Switzerland; Brain institute and Cancer Center, UT Southwestern Medical Center, 75235-8823 Dallas, Texas, United States; Department of Anatomy and Neurosciences, section Clinical Neuroanatomy and Biobanking, Amsterdam Neuroscience, Amsterdam University Medical Centre, Vrije University Amsterdam, 1081 HZ, Amsterdam, The Netherlands; Amsterdam Neuroscience, program Neurodegeneration, Amsterdam University Medical Centre, Vrije University Amsterdam, 1081 HZ, Amsterdam, The Netherlands; School of Life Sciences, Ecole Polytechnique Fédérale de Lausanne, 1015 Lausanne, Switzerland

## Abstract

Correlative light and electron microscopy (CLEM) combines light microscopy, for identifying a target via genetic labels, dyes, antibodies, and morphological features, with electron microscopy, for analyzing high-resolution subcellular ultrastructures. Here, we describe the step-by-step instructions to perform a CLEM experiment, optimized for the investigation of ultrastructural features in human brain tissue. The procedure is carried out at room-temperature and can be also adapted to other human and animal tissue samples. The procedure requires 8-days to complete and includes the stages of sample fixation for optimal ultrastructural preservation, immunofluorescence staining, image acquisition, multi-modal image correlation, and is executable within standard EM laboratories. Serving as a critical tool for characterizing human tissue and disease models, room-temperature CLEM facilitates the identification and quantification of subcellular morphological features across brain regions.

**Key points:**

- The protocol for correlative light and electron microscopy (CLEM) is optimized for analyzing chemically fixed human brain tissues. It focuses on maintaining the integrity of ultrastructural features, thereby minimizing artifacts and structural alterations.
- Examining brain tissues at the ultrastructural level can provide an unprecedented amount of detail which may help advance our understanding of the mechanisms underlying neurodegenerative disorders.

## Introduction

Room temperature electron microscopy (EM) is a powerful tool to investigate biological samples label-free at high resolution down to the nanometer scale, which allows the visualization of sub-cellular features. collectively called “ultrastructure”. However, EM is limited by its field of view. Much larger areas and volumes can be investigated using light microscopy (LM) although conventional LM is limited by its resolution, roughly at 250 nm due to the diffraction limit of light. LM predominantly utilizes dyes or fluorescent labeling to selectively illuminate cellular components, constraining the analysis to the spectral capabilities of the microscope. In comparison, the resolution offered by EM remains unmatched, allowing for visualization of ultrastructural features at a finer scale.

The integration of LM and EM in correlative light and electron microscopy (CLEM) enables the scanning of large fields of view with LM and targeted investigation of regions of interest (ROIs) by EM. This approach is particularly beneficial for studying rare targets or when specific biochemical characterizations using markers are required, offering significant advantages over using EM alone.

CLEM requires meticulous methodology and specific technical expertise. Successful CLEM depends on carefully balancing the preservation of ultrastructure, which is prone to quick deterioration, with effective LM staining that highlights specific targets. For the field of neurodegeneration, working with human brain tissue presents additional challenges, notably the lack of genetically encoded molecular labels and the risk of ultrastructural damage caused by post-mortem delay and during permeabilization for immunolabeling^1^. However, advancements have significantly enhanced the capability of CLEM. These include the development of high-affinity antibodies for specific epitopes and posttranslational modifications, which improve specificity and sensitivity. Improved EM preparation methodologies have also extended tissue viability for more detailed analysis^2–5^. Enhancements in labeling strategies^6^, super-resolution fluorescence microscopy^7^, faster and higher-resolution electron detectors, and sophisticated image analysis algorithms^8^ enable more detailed exploration of complex biological processes and disease mechanisms at the nanometer scale. These technological improvements are vital for advancing our understanding of biological structures and functions.

Our protocol is straightforward and adaptable, designed by a multidisciplinary team with extensive expertise in advanced microscopic imaging for studying chemically fixed human brain. It has been effectively used to analyze protein aggregates in various neurodegenerative diseases such as Parkinson’s disease,^9,10^, multiple system atrophy^11^ and frontotemporal lobar dementia^12^. Additionally, the protocol has been applied to cell culture samples^12,13^, human skin biopsies and mouse tissues (^14^ and Supplementary Figure 1). The basic protocol only requires minimal specialized equipment, allowing anyone with access to a light microscope, an ultramicrotome and an electron microscope to perform this CLEM pipeline independently.

### Development of the protocol

For the study of chemically fixed human brain, we developed a post-embedding CLEM approach (Figure 1) to examine the ultrastructure of Lewy bodies in Parkinson’s disease^9^. We collected human brain tissue within six hours post-mortem delay, followed by chemical fixation to maximize the ultrastructural preservation of the tissue for EM. Vibratome sections (20-100 micron thick) were heavy metal stained and flat embedded in resin. Small pieces (∼1 × 1 mm) were mounted on resin blocks for serial sectioning. Serial sections were collected alternately on glass slides and EM grids. The slides were processed for immunohistochemistry (IHC) with ROIs localized by positive antibody staining detected by LM. Adjacent EM grids were imaged by transmission EM (TEM) at low magnification to map the ROIs. LM and EM images were overlaid to pinpoint the location of the ROIs in the TEM images for higher resolution imaging.

**Figure 1.**
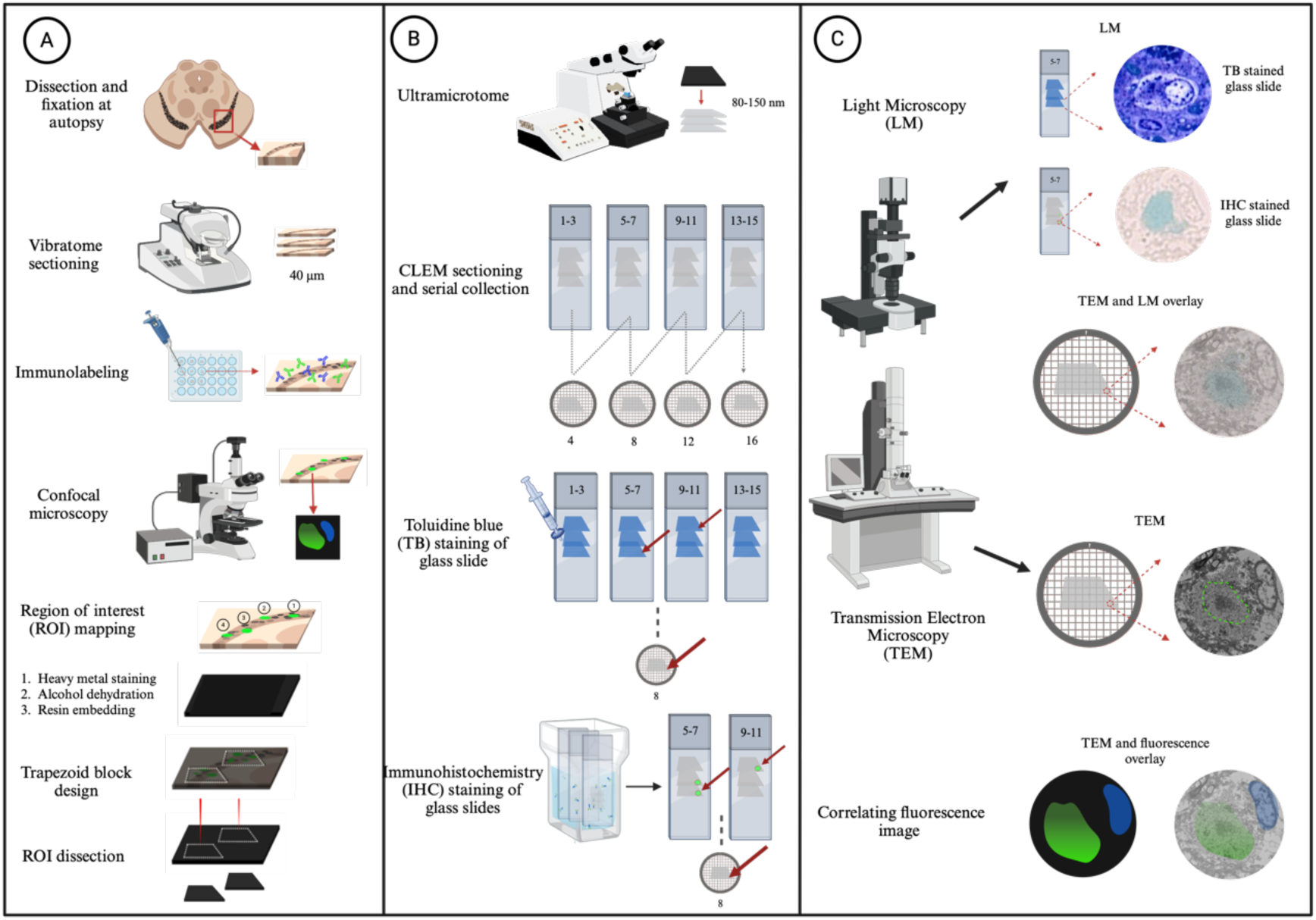
Schematic of the CLEM workflow; A) Brain samples are dissected and fixed at autopsy in 0.1% glutaraldehyde and 4% paraformaldehyde before being sectioned into 40 µm sections with a vibratome. The free-floating tissue sections are immunolabeled and imaged with fluorescence-based microscopy, computationally annotating and mapping the ROIs onto a section overview of the image. The imaged section is prepared for EM through *en bloc* staining and resin embedding. Using the ROI map as guidance, trapezoid blocks containing ROIs are excised from the resin-embedded tissue sections B) The excised tissue blocks are sectioned into 80-150 nm ultrathin sections which are serially collected on glass slides and EM grids. Toluidine blue (TB) and/ immunohistochemistry (IHC) staining is performed on the glass slides to identify the exact adjacent grid containing the ROI; C) The light microscopy (LM) images are used to guide the collection of high-resolution EM images and act as an intermediate bridge to correlate the fluorescence microscopy with the EM ultrastructure.

We subsequently enhanced the efficiency of our protocol by immunolabeling vibratome sections, followed by fluorescent-based microscopy allowing the targeting of tissue areas with abundant pathology and a detailed 3D fluorescent characterization of specific ROIs^10,11^. Toluidine blue staining of the LM aided in correlating fluorescence, LM, and EM images^15^, and laser cutting enabled the precise excision of ROIs based on the fluorescence signals. These improvements have significantly enhanced our CLEM pipeline, enabling accurate targeting and analysis of regions with sub-cellular precision in the human brain ^10,11^.

### Applications of the method

The CLEM approach presented here has primarily been used to investigate neuropathological aggregates in post-mortem brains of neurodegenerative disease patients^9–12^. We have also applied it to human skin biopsies (Supplementary Figure 1), mouse tissues (^14^ and Supplementary Figure 1) and on cell monolayers^12,13^, demonstrating its versatility for a wide range of potential applications in biology. This step-by-step protocol for chemically fixed human brain tissue can be used in any EM laboratory and can be also coupled to correlative volume EM approaches such as serial block face scanning electron microscopy (SBF-SEM) or focused ion beam scanning electron microscopy (FIB-SEM)^16^ and array tomography, if the high resolution of TEM is not required^17–19^.

### Comparison with other methods

Several notable and innovative room-temperature CLEM protocols have recently been developed. For example, super-resolution fluorescence CLEM for cell culture samples^20^, utilizing a two-photon laser microscope to study synapses in the mouse brain^21^, dark field microscopy combined with SEM to track endocytosis and the intracellular distribution of spherical nucleic acids^22^ and fluorescence recovery after photobleaching microscopy paired with EM to examine labeled vesicles in synapses of cultured hippocampal neurons^23^. Each of these protocols are tailored to specific sample types and incorporate distinct technical methods. Our protocol employs standard EM and LM techniques and equipment and, while primarily focused on the human brain, is versatile enough to be adapted for studying other human and animal tissues, as well as cell monolayers^12,13^. We provide detailed guidance on how to preserve tissue ultrastructure and facilitate the correlation between fluorescence, LM and EM images.

### Experimental design

This protocol describes how to investigate the ultrastructure of chemically-fixed human brain tissue using CLEM. The procedure is illustrated in Figure 1. We also included details where the protocol would differ significantly for the investigation of other human and mouse tissues. We used openly available software plugins, developed custom-scripts and used specific instrumentation to maximize precision and automation where possible. However, these automation steps are not essential to the protocol and can be done manually. We detail these additions as optional.

### Sample preparation

#### Tissue fixation

We use post-mortem human brain tissue collected using a rapid autopsy protocol with less than 6 hours post-mortem delay to limit the effects of tissue decay on the ultrastructure^1^. The tissue is ideally fixed at autopsy in 4% paraformaldehyde (freshly dissolved in buffer, or commercially available) and 0.1% glutaraldehyde in 0.15 M cacodylate buffer for 24 hours. Cacodylate buffer is preferred for its reported ability to neutralize free radicals in neural tissue^24–26^. However, due to its high toxicity and banning in many countries, it can be replaced by phosphate buffer pH 7.4 for fixation^27^. According to our testing this is the optimal chemical fixative for preserving the tissue ultrastructure whilst retaining some tissue antigenicity^1^. Prospective rapid autopsy collection is not always feasible, but good tissue ultrastructure can be preserved by post-fixation of paraformaldehyde-only fixed samples. Optimal ultrastructure will be maintained if post-fixation in glutaraldehyde is carried out within a year of autopsy, with integrity declining over time in buffered formalin (formaldehyde and methanol in water) fixation.

#### Antibody labeling

The brain blocks are vibratome sectioned and stained with fluorescent dyes and antibodies to localize regions of interest, avoiding permeabilization or antigen retrieval methods (e.g. detergents, picric acid, heat, formic acid) to maintain ultrastructural integrity. Chemical fixation creates holes in the membranes that allow many antibodies to penetrate, however glutaraldehyde can negatively impact the labeling and cause autofluorescence, so appropriate imaging controls (e.g. no primary antibody) should be included. Multi-labelling can tag the protein/structure of interest and the surrounding cellular features which could be used later as fiducial markers for image correlation. The number of labels that can be utilized is limited only by the number of channels available on the fluorescent microscope. DAPI penetrates glutaraldehyde-fixed tissue well up to at least 200 µm thickness, and we have characterized several cell-specific markers (^11^ and Supplementary Table 1). The antibody concentration, staining time and tissue thickness should be optimized for each antibody used.

#### Fluorescent imaging

The free-floating sections are reversibly mounted on glass slides and imaged using fluorescence microscopy. The fluorescent labeling is used for mapping all ROIs in a given brain section. High resolution and volumetric z-stacks of the individual ROIs can be recorded here as well. Our protocol describes the imaging of the sample by widefield fluorescence microscopy, however confocal or super-resolution microscopy (STORM, PALM, etc.) could also be used depending on the resolution and fluorescent characterization of ROIs required. Bleaching of the sample is not a limitation in this protocol as the fluorescence is not maintained after osmium impregnation and is not detected on EM sections.

#### En bloc staining and resin embedding

After finding and mapping the ROIs, the section is stained with heavy metals, dehydrated and embedded in resin, for TEM imaging. The *en bloc* staining protocol follows published protocols (rOTO + uranyl acetate + lead aspartate) without any significant modifications^2,3,5^. Due to strict regulations on uranyl acetate, a good alternative could be the use of neodymium acetate^28^. A high contrast protocol (for SBF-SEM and FIB-SEM applications) and a shorter and more classic protocol (reduced Osmium - Osmium or only reduced Osmium) for TEM only is provided.

We describe the manual *en bloc* staining and resin embedding protocol but have also optimized it for an automated tissue processor. For the latter, we custom-designed a sample holder for 40-150 micron vibratome sections to minimize excessive movement and curling/folding (Supplementary Figure 2). This holder can be built in-house with a 3D printer (however professional machining in surgical metal is recommended) using the provided schematics and material information (Supplementary Figures 3-4, Supplementary Data 1 and 2). In all cases, flat embedding is required to be able to image the sections by LM.

### Preparation of blocks for CLEM sectioning

The regions in the sample containing the ROIs must be excised out of the flat embedded sample and glued on an empty resin block in preparation for serial sectioning. The embedded sections are first imaged with brightfield microscopy and then overlaid with the previously acquired fluorescence maps to mark the areas which need to be excised. The BigWarp^29^ plugin in FIJI^30^ can be used for a more precise alignment of the two images. The blocks can be excised manually with a razor blade or scalpel using macro-features in the tissue (section edges, arteries, large myelin tracts, different tissue densities etc.) for accuracy. Such features will be more easily visible if the short staining protocol is used. Alternatively, we use the aid of laser microdissection to increase the precision and efficiency of this step. The details of this protocol are provided in the Supplementary Information. The area of the block that is to be excised is up to the user’s discretion, however, must be maximally limited in the X and Y dimensions to the area that will fit on the EM grids.

### CLEM sectioning and immunohistochemistry (IHC)

In preparation for CLEM sectioning, the excised tissue is mounted on resin blocks. An ultramicrotome is used to sequentially cut ultra-thin (<100 nm) and semi-thin (200 nm) sections, collected alternatively on EM grids and glass slides, respectively. Glass slides stained with toluidine blue dye provide high contrast and morphological details comparable to low magnification TEM^15^. The slides are imaged by brightfield microscopy to locate the ROI and follow the tissue morphology for later correlation with fluorescence and/or EM. Optionally, immunohistochemistry staining (IHC) can be performed on the glass slides where antibodies, visualized with chromogenic dyes (e.g. DAB, or similar), can target proteins of interest with the aim of identifying protein aggregates, cell types, or pathological features. Antigen retrieval during this IHC step allows for the use of additional antibodies, that were not able to be included in the fluorescence, here instead. Additionally, multiplexed labeling can be utilized. The stained slides are imaged by LM to locate the ROI within the sections and to follow the morphology of the tissue for later correlation with fluorescence and/or EM.

The alternate collection of serial sections on glass slides and EM grids is a key step in this technique, as it enables relocalization of the ROI within the embedded volume, acting as an intermediate bridge for the efficient correlation of fluorescence volume with the individual 2D ultrathin LM/EM images. The IHC step is useful for identifying which EM grid contains the ROI, particularly when it is difficult to detect in the monochromatic landscape of the toluidine blue image, or if fluorescence pre-localization has not been performed.

### EM Imaging and Correlation

Bright field images of IHC sections are used to identify which cycles (a set of adjacent serial sections collected on glass slide and EM grids) contain immunopositive ROIs. By comparing tissue features and fiducials visible on both EM images and bright field images of the same cycle, ROIs are localized on EM grids. To facilitate this, a Python script (https://github.com/LBEM-CH/clem4cina) was developed to correlate XY coordinates on LM images with those on the EM grid (Supplementary Information). This software is especially useful for difficult localization of ROIs in crowded ultrastructural landscapes, such as brain tissue.

Correlating fluorescent microscopy and EM images is challenging due to their size difference (two orders of magnitude in width and length, three orders of magnitude in depth). Toluidine blue sections, collected adjacent to IHC sections, show visible tissue features (e.g. cell nuclei, myelin sheaths and blood vessels) that serve as landmarks for image correlation. We used the interactive FIJI-plugin BigWarp^29^ to align images based on user-defined landmarks. However other correlation software could also be used (e.g. eC-CLEM^31^).

Post-processing of a TEM tilt-series, such as tomography and segmentation, can provide 3D information. Additionally, if the serial sectioning step is omitted, the same preparation protocol can be used to prepare samples for volume EM methods such as SBF-SEM or FIB-SEM.

### Expertise needed to implement the protocol

Our protocol assumes an understanding of basic wet-lab procedures and the concept of immunolabeling. In addition, researchers need to be experienced with vibratomes and ultramicrotomes for sample preparation, as well as brightfield, fluorescence, and electron microscopy for imaging. Users need to follow the safety rules and the regulations of their respective facility center when handling human brain tissue and all chemicals used in this protocol. Some familiarity with FIJI is helpful, but the publicly available software documentation should be sufficient to guide users through this software for image correlation.

### Limitations

We have used this CLEM pipeline to investigate subcellular ultrastructure within post-mortem human brain tissue^9–13,32^, other human and mouse tissues (^14^ and Supplementary Figure 1) as well as cell monolayers^12,13^. However, this protocol has several limitations that need to be considered. First, the native state of the sample will be altered because of chemical fixation, heavy metal staining and resin embedding. The specific changes due to these treatments have been well-documented elsewhere and will not be discussed here^1,3,33–36^. The heavy metal staining, used for EM, limits the achievable resolution to approximately 2 nm, making it unsuitable for atomic structural analysis of proteins. EM sections are maximally limited to 200 nm, restricting large volumetric information, and the field of view is constrained to the EM grid size (the maximum diameter is 3 mm). The correlation of ROIs for the adjacent sections of IHC glass slides and EM grids is limited by the cutting thickness. Following this protocol, the minimum distance between ROIs on glass slides and EM grids is 150 nm ((200 nm semi-thin LM section + 100 nm consecutive ultra-thin EM section)/2) which represents the maximum achievable correlation in the z-plane. However, if thinner sections are cut smaller features can be correlated. Furthermore, limitations may arise with antibody usability in EM fixed tissue, as well as the antibody penetration depth^11,37^. In the event that neither the protocols we provide here for fluorescence labelling or on-resin IHC are effective, methods such as pre-embedding labelling using ultra-small gold conjugates could prove effective^38^. Overall, the protocol is time-consuming and labor-intensive, and therefore unsuitable for high-throughput EM analysis. However, depending on the ROI distribution in the sample, it may be possible to obtain all the information necessary from only a few samples.

### Regulatory approvals

Approvals may be required to obtain bio-specimens. The timeline and costs will be dependent on the specific regulations imposed by the organizations involved. All brain samples used for this study were prospectively collected from participants in the Netherlands Brain Bank donation program (www.brainbank.nl). The NBB adheres to strict ethical guidelines such as written informed consent, open access, and non-profit policy, as stated in BrainNet Europe’s Ethical Code of Conduct for brain banking. The procedures of the NBB are in compliance with Dutch and European law and have been approved by the Ethics Committee of the VU University Medical Center in Amsterdam, the Netherlands. Approval of the scientific advisory board and ethics committee for the prospective collection of postmortem brain tissue for the current project was received within eight weeks. The dissection and processing of the postmortem brain tissue was performed by WvB (AmsterdamUMC). A Material Transfer Agreement (MTA) was signed to ensure that conditions for the transfer of material and data are agreed by all parties (see for more details: Information for tissue applicants_2019.pdf).

## Materials

### Reagents

#### Sample Fixation, vibratome sectioning and fluorescence labeling

- Paraformaldehyde 16% aqueous solution EM grade (Electron Microscopy Sciences, cat. no. 15700) CAUTION-Paraformaldehyde is harmful if swallowed or inhaled. It may irritate the skin. Wear suitable clothing, gloves and eye protection and work under the fume hood.
- Glutaraldehyde 25% aqueous solution EM grade (Electron Microscopy Sciences, cat. no. 16200) CAUTION-Glutaraldehyde irritates the skin. Wear suitable clothing and gloves and work under the fume hood.
- Sodium cacodylate (Sigma-Aldrich, cat. no. C0250) CAUTION-Sodium cacodylate is irritant and toxic if swallowed. Wear suitable clothing and gloves, and work under the fume hood. This chemical is banned by many countries and institutions. It can be replaced by phosphate buffer for fixation
- Glycerol (Sigma-Aldrich, cat. no. 1.04095)
- Hydrochloric acid (HCl, sigma-Aldrich, cat. no. 1.00317) CAUTION-Hydrochloric acid burns the skin and damages the eyes. Wear suitable clothing, gloves, eye protection and work in a fume hood.

#### *En bloc* staining and resin embedding

- Distilled deionised H2O (ddH2O)
- Calcium chloride (Sigma-Aldrich, cat. no. C4901)
- Acetone 99.5% (Acetone, Sigma Aldrich, cat. no. 32201-1L) CAUTION-Acetone is flammable and toxic. Keep them in an allocated cupboard. Wear suitable clothing and gloves.
- Ethanol (Ethanol, Fischer Chemical, cat. no. E/0650DF/15) CAUTION-Ethanol is flammable and toxic. Keep them in an allocated cupboard. Wear suitable clothing and gloves.
- L-aspartic acid (Sigma-Aldrich, cat. no. 11189)
- Lead nitrate (Sigma-Aldrich, cat. no. 228621) CAUTION-Do not breathe the lead nitrate dust and avoid contact with skin or eyes. Wear suitable clothing and gloves and work in a fume hood.
- Osmium tetroxide 4% Aqueous solution (Electron Microscopy Sciences, cat. no. 19160) CAUTION-Osmium tetroxide is toxic if inhaled, comes into contact with skin and if swallowed. Wear suitable clothing, gloves and eye protection and work in a fume hood.
- Potassium ferrocyanide (Sigma-Aldrich, cat. no. P3289) CAUTION-Potassium ferrocyanide is irritant. Wear suitable clothing and gloves.
- Durcupan resin: supplied in four separate parts (Sigma-Aldrich, Ducupan Resin A/M, cat. no. 44611; Sigma-Aldrich, single component B, hardener 964, cat. no. 44612; Sigma-Aldrich, single component D, cat. no. 44614; Electron Microscopy Sciences, DMP-30 cat. no. 13600) CAUTION-Durcupan resin is toxic by inhalation, in contact with skin and if swallowed. Wear suitable clothing, gloves and eye protection and work under the fume hood.
- Sodium cacodylate (Sigma-Aldrich, cat. no. C0250) CAUTION-Sodium cacodylate is irritant and toxic if swallowed. Wear suitable clothing and gloves.
- Sodium hydroxide (NaOH, Sigma-Aldrich, cat. no. 221465) CAUTION-Sodium hydroxide causes severe burns. Avoid contact with skin and eyes. Wear suitable clothing, gloves and eye protection.
- Thiocarbohydrazide (Sigma-Aldrich, cat. no. 223220) CAUTION-Thiocarbohydrazide is toxic if inhaled, comes into contact with eyes and skin and if swallowed. Wear suitable clothing, gloves and eye protection and work under the fume hood.
- Uranyl Acetate (Electron Microscopy Sciences, cat. no 22400) CAUTION-Uranyl Acetate is radioactive material and heavy metal. When dealing with radioactive material, appropriate safety precautions must be followed. Do not breathe the uranyl acetate dust and avoid contact with skin or eyes. Wear suitable clothing and gloves and work in a fume hood

#### Serial sectioning

- Distilled deionised H2O (ddH2O)
- Ethanol (Fischer Chemical, cat. no. E/0650DF/15) CAUTION-Ethanol and acetone are flammable and toxic. Keep them in an allocated cupboard. Wear suitable clothing and gloves.
- Borax (Electron Microscopy Sciences, cat. no. 21130)
- Toluidine Blue (Electron Microscopy Sciences, cat. no. 22050) CAUTION-Toluidine Blue is toxic if inhaled. Wear suitable clothing, gloves and eye protection.

#### Immunohistochemistry (IHC)

- Distilled deionised H2O (ddH2O)
- Potassium hydroxide (KOH, Sigma-Aldrich, cat. no. 1.05033.0500) CAUTION-Potassium hydroxide irritates and burns the skin. It is toxic if swallowed. Wear suitable clothing, gloves and eye protection.
- Triton X-100 (Sigma-Aldrich, cat. no. T9284) CAUTION-Triton X-100 is harmful when it comes into contact with the eyes and skin. Wear suitable clothing, gloves and eye protection.
- Hydrogen peroxide (H2O2, Sigma-Aldrich, cat. no. 1.07210.0250) CAUTION-Hydrogen peroxide can cause irritation to the eyes and skin. Wear suitable clothing, gloves and eye protection
- Methanol 100% (Sigma-Aldrich, cat. no. 34860-1L-R) CAUTION-Methanol is flammable and toxic. Keep them in an allocated cupboard. Wear suitable clothing and gloves
- Ethanol 100% (Fischer Chemical, cat. no. E/0650DF/15) CAUTION-Ethanol is flammable and toxic. Keep them in an allocated cupboard. Wear suitable clothing and gloves
- Antibody diluent (Agilent Dako REAL, cat. no. S202230), or suitable alternative
- Primary antibody (specific to protein of interest)
- Secondary antibody (according to the primary antibody species. Anti-mouse, Reactolab SA, cat. no.MPX-2402-15; Anti-rabbit, Reactolab SA, cat. no. MP-7401-15; Anti-goat, Reactolab SA, cat. no. MP-7405-15)
- HRP-green chromogen kit (Zytomed Systems, cat. no. ZUCO70-100)
- Mounting solution, NeoMount (Sigma-Aldrich, cat. no. 109016)
- Sodium chloride (NaCl - Fisher Chemical, cat. no. S/3160/60)
- Disodium hydrogen phosphate (Na2HPO, Sigma-Aldrich, cat. no. S9763)
- Potassium chloride (KCl, Roth, cat. no. 6781.1)
- Potassium phosphate monobasic (KH2PO4, Sigma-Aldrich, cat. no. P0662) CAUTION-Do not breathe the potassium phosphate monobasic dust and avoid contact with skin or eyes. Wear suitable clothing and gloves
- (OPTIONAL) Formic acid (Sigma-Aldrich, cat. no. F0507) CAUTION-When using formic acid, it should be performed under a fume hood due to its irritants and burning effects on the skin and eyes
- (OPTIONAL) Tris (Sigma-Aldrich, cat. no. 93352)
- (OPTIONAL) Ethylenediaminetetraacetic acid (EDTA, PanReac AppliChem, cat. no. A5097,0500)
- (OPTIONAL) Sodium citrate (Sigma-Aldrich, cat. no. 71402-250g)
- (OPTIONAL) Citric acid (Sigma-Aldrich,cat. no. C1909) CAUTION-Citric acid is irritant. Wear suitable clothing and gloves

### Reagent Setup

#### 3 mM calcium chloride solution

- Dissolve 1.1 g of calcium chloride to 10.0 ml of ddH2O.

#### 0.3 M cacodylate buffer

- Dissolve 12.8 g of sodium cacodylate to 180.0 mL of ddH2O and adjust the pH to 7.4 with HCl/NaOH, supplement with 3.0 mM of calcium chloride (0.6 mL) and make up the final volume to 0.2 L with ddH2O. CRITICAL-Sodium cacodylate is optimal for the preservation of membrane ultrastructure in neural tissue. Phosphate-based buffers can be used instead for fixation however rinsing steps have to be carefully performed to avoid precipitates 4% paraformaldehyde, 0.1% glutaraldehyde (v/v) in 0.15 M cacodylate buffer as EM fixation buffer:
- Mix 2.5 mL of 16% paraformaldehyde and 40.0 µL of 25% glutaraldehyde with 5.0 mL of 0.3 M cacodylate buffer and make up the final volume to 10.0 mL with ddH2O.

#### 0.1% (v/v) paraformaldehyde in 0.15 M cacodylate buffer as storage buffer

- Mix 62.5 µL of 16% paraformaldehyde with 5.0 mL of 0.3 M cacodylate buffer and make up the final volume to 10.0 mL with ddH2O.

#### 10 M tris-buffered saline (TBS)

- Dissolve 24.0 g of tris base and 88.0 g of NaCl in 0.9 L ddH2O water. Adjust pH to 7.4 with 1.0 M HCl and and make up the final volume to 1.0 L with ddH2O.

#### 50% (v/v) glycerol solution in TBS for fluorescence imaging

- Mix 1.0 ml of Glycerol with 1.0 ml TBS buffer.

CRITICAL-glycerol at a slow rate due to its viscosity.

### Aspartic acid for *en bloc* staining and resin embedding the tissue

- Weigh out 0.2 g aspartic acid and add to 50.0 mL ddH2O. Leave in a 60°C oven overnight to dissolve completely before use. The volume of solution prepared should be prepared 2-3 times in excess of what is needed.

### 2% (v/v) aqueous osmium tetroxide for *en bloc* staining and resin embedding the tissue

- Add 5.0 mL of 4% aqueous osmium tetroxide to 5.0 mL ddH2O.

### Graded alcohol series (20%, 50%, 70%, 90%, 100%)

- Prepare 10.0 mL each of the indicated percentages (v/v) of ethanol or acetone in ddH_2_O. CRITICAL-If using a well-plate make sure the plastic is resistant to acetone or use ethanol for dehydration.

### 2% (v/v) reduced osmium for *en bloc* staining and resin embedding the tissue

- Add 2.0 mL of 0.3 M cacodylate buffer to 0.15 g of potassium ferrocyanide powder into a weight boat and pour into a 15.0 mL conical tube. Vortex well to dissolve before adding 5.0 mL of 4% aqueous osmium. Top up to a total volume of 10.0 mL with 0.3 M cacodylate buffer. The final concentration of the solutions should be 2% (v/v) osmium tetroxide, 0.15 M cacodylate buffer and 0.15% (w/v) potassium ferrocyanide.

### Durcupan resin for *en bloc* staining and resin embedding the tissue

- Weigh out 10.0 g each part A and B with 0.3 g part D into a plastic wide mouth bottle. Mix thoroughly by stirring slowly on a magnetic stirrer. Add 0.8 ml of DMP-30 and mix thoroughly for another 10 minutes. The mixture turns red upon adding DMP-30. Keep stirring slowly until use but try to avoid air bubble formation.

CRITICAL-Resin should be prepared fresh immediately before use.

CAUTION-Resin should be prepared and used in a fume hood at all times. Excess resin should be polymerized in the wide mouth bottle in a 60°C oven for 48 hours before discarding.

### 1.0 M sodium hydroxide (NaOH)

- Add 2.0 g of NaOH to 50.0 mL ddH2O and agitate to dissolve.

### 1% (w/v) thiocarbohydrazide (TCH) for *en bloc* staining and resin embedding the tissue

- Add 0.1 g thiocarbohydrazide to 10.0 mL ddH2O and place in a 60°C oven for 1 hour, mixing by inversion every 10 minutes. Cool down to room temperature and filter with a 0.22 µm syringe filter before using. CRITICAL-Some batches of TCH can produce a purplish-black solution after dissolving at 60°C, rather than being colourless. However, this does not affect the samples.

### 2% (w/v) uranyl Acetate for *en bloc* staining and resin embedding the tissue

- Add 0.2 g of uranyl acetate powder to 10.0 mL ddH2O and agitate gently. Leave at room temperature to dissolve completely. Filter through a 0.22 µm syringe filter before use. CAUTION-Uranyl acetate is radioactive. Follow internal safety rules for radioactive materials.

### Waltons lead aspartate for *en bloc* staining and resin embedding the tissue

- Add 0.066 g lead nitrate per 10.0 mL of 60°C aspartic acid and mix by inversion. Keep the solution in a prepared 60°C water bath. Adjust pH to 5.5 (at 60°C) dropwise with newly prepared 1.0 M NaOH. Stabilize at 60°C for 30 mins. CRITICAL-This step is prone to precipitation if the temperature drops below 60°C, and/or if the NaOH is not fresh enough. The pH of the solution should be adjusted while the temperature is being maintained in the 60°C water bath. The pH of the solution changes quickly above pH 4.0 so the NaOH should be added dropwise with thorough stirring/swirling in between. In case the pH of the solution reaches above 5.5, it should not be corrected by the addition of acid. The solution must be prepared again from the beginning using the excess 60°C aspartic acid.

### Toluidine blue solution

- Dissolve 0.1 g borax in 10.0 mL ddH2O. Add 0.1 g toluidine blue powder in the borax solution. NOTE – Borax is added to the toluidine blue solution to help the stain penetrate epoxy resins^39^.

CRITICAL-Store in a 10.0 mL syringe with a 0.22 µm sterile syringe filter at room temperature (RT). CAUTION-Have a dedicated waste container for the disposal of toluidine blue solution in accordance with internal regulations for the disposal of hazardous chemical waste. It is a strong dye, use ethanol to clean surfaces if spilled.

### Saturated potassium hydroxide (KOH) using as etching solution

- Stir 11.0 g KOH in 50.0 mL ethanol at RT overnight.

### 1x PBS, pH 7.4

- Mix 80.1 g NaCl, 14.4 g Na2HPO4, 2.0 g KCl and 2.7 g KH2PO4 to 0.8 L ddH20. Adjust pH to 7.4 and make up the final volume to 1.0 L with ddH2O.

### 20% (v/v) Triton X-100 in PBS (PBS-T)

- Stir 0.5 mL of Triton X-100 in 0.2 L PBS for at least 1 h. CRITICAL-Pipette Triton-X 100 at a slow rate due to its viscosity.

### (Optional) 1x tris-EDTA

- Mix 1.2 g tris, 0.37 g EDTA to 0.8 L ddH2O until dissolved. Adjust pH to 9.0 and make up the final volume to 1.0 L with ddH2O.

### (Optional) 1x sodium citrate

- Mix 24.3 g sodium citrate dihydrate and 3.4 g of citric acid to 0.8 L ddH2O until dissolved. Adjust pH to 6.0 and make up the final volume to 1.0 L with ddH2O.

### Equipment

#### Sample fixation, vibratome sectioning and fluorescence labeling

- Vibratome (Leica VT1200S)
- Cyanoacrylate glue (Biosystems AG, cat.no 14037127414)
- Narrow-tip paint brushes (Electron Microscopy Sciences, cat. no. 65575-02)
- Double-sided razor blades (Science Services, cat. no. E72000)
- Light microscope equipped with a camera to detect and image-stained ultra-thin tissue sections (Leica THUNDER Imager 3D Tissue equipped with Flexacam C1 camera)
- Fluorescent microscope (Leica THUNDER Imager 3D Tissue equipped with a K8 fluorescent camera)
- (Optional) Laser cutting machine capable of importing coordinates for precision cutting (Leica LMD7)
- Glass slides (VWR, cat. no. 630-1985)
- Precision cover glasses (Marienfeld, cat. no. 0107222)
- Microscope coverslips (Epredia, cat. no. BB02400240A153FST0)

### *En bloc* staining and resin embedding

- Magnetic stirrer and stirring bars
- Orbital shaker (Labnet Orbit LS), vial rotator (Pelco R2 Rotary mixer) or tissue processor (Leica EM TP)
- Oven (Capable to reach 60°C)
- pH meter (with temperature probe highly advantageous)
- Water bath
- Weighing scales
- Aclar 33C Embedding Film (Electron Microscopy Sciences, cat. no. 50425-10)
- 15 and 50 mL centrifuge tubes (ThermoFisher Scientific, cat. no. 33.96.50; cat. no. 339652)
- Flat embedding molds (14mm x 5mm x 3mm, Electron Microscopy Sciences, cat. no. 70902)
- Glass scintillation vials with a poly-seal cone liner (Electron Microscopy Sciences, cat. no. 72632) or multi-well plates (12 wells, polystyrene plate, flat bottom. Sigma-Aldrich, cat. no. CLS3512)
- Tooth-picks / applicator sticks (Electron Microscopy Sciences, cat. no. 72304-10)
- 20 ml, 10 ml and 5 ml syringes (VWR, cat. no. 720-2521; cat. no. 720-2520; cat. no. 720-2519)
- Sterile syringe filters 0.22 µm (Merck, cat. no. SLGV013SL)
- Plastic wide mouth bottle (Fisherbrand, cat no.02-896B)

### Serial sectioning

- Ultramicrotome (Leica, EM UC7)
- Diamond knife (Ultra 45°, Diatome, cat. no. DU4515) CAUTION-It is extremely important to keep diamond knives in a safe place. For the user and knife’s safety, it is important not to touch the knife’s tip.
- Trimming knife (Trim 90, Diatome, cat. no. DTB90) CAUTION-It is extremely important to keep diamond knives in a safe place. For the user and knife’s safety, it is important not to touch the knife’s tip.
- Hot plate (Labnet, Accuplate Hotplate Stirrer) CAUTION-Avoid touching the hot plate while it is on.
- 2 x Eyelash manipulator (Electron Microscopy Sciences, cat. no. 71182)
- Double-sided razor blades (Science Services, cat. no. E72000) CAUTION-Razor blade is sharp. Avoid touching the edge of the razor blade.
- Electron microscopy grids (Electron Microscopy Sciences, cat. no. FF-100H-Cu-UL, cat. no. FCF2010-CU-TA-50ST)
- Grid storage box (Electron Microscopy Sciences, cat. no. 71152)
- Superfrost plus glass slides (BioSystems, cat. no. 85-0915-00)
- Qualitative filter paper (Whatman, cat. no. 1001090)
- Diamond scribing tool (Electron Microscopy Sciences, cat. no. 70030)
- Super glue (Loctite Adhesive 410, Digitec Galaxus, cat. no. 6489185)
- Reverse action tweezer (Ideal-tek, cat. no. 3X.SA.1)
- Resin stubs previously prepared
- Loops (Electron Microscopy Sciences, cat. no. 70944)

### Immunohistochemistry (IHC)

- Steamer (Tristar steamer 12 L)
- Oven (Capable to reach 37°C)
- PAP pen (Agilent, cat. no. S2002002230-2)
- Staining tray (Fisher Scientific, cat. no. 155189960)
- Staining jars (Fisher Scientific Kartell, cat. no. 10375681)
- Glass staining jar (VWR, cat. no. 631-0698)

### Other

- Transmission electron microscopy 80 kV-120 kV (Tecnai G2 Spirit 80kV, ThermoFisher; T12 120 kV, FEI)
- Computer with a 64-bit operating system and system requirements capable of supporting the choice of correlation software (For FIJI: Windows XP, Vista, 7, 8, 10, 11; Mac OS X 10.8 “Mountain Lion” or later; Linux on amd64 and x86 architectures).
- Correlation software (FIJI^30^)
- Fume hood

### Procedure

#### Sample fixation Timing: 2.5 – 24.5 hours depending on the sample

1. Fix the samples for EM according to the protocol for (a) human brain (24.5 hours) or (b) mouse tissues (2.5 hours).

a. Sample fixation for human brain:

i. Dissect brain or tissue blocks based on anatomical landmarks, volume 0.5 cm^3^.
ii. Immerse the brain blocks in EM fixation solution for 24 hours at 4°C.
iii. Transfer the samples into the storage solution and keep at 4°C. NOTE-If the tissue blocks were not fixed for EM at autopsy, post-fix the tissue blocks in EM fixative, ON, 4°C.
b. Sample fixation for mouse tissues:

i. Anesthetize the mouse following local institutional guidelines. Perfuse the mouse via the heart with a perfusion pump in EM fixation solution at a speed of 7 ml/minute using approximately 250-300 ml per animal.
ii. After perfusion, leave the animal for 1 h.
iii. Dissect the relevant tissues according to local institutional guidelines.
iv. Transfer the samples into the storage solution and keep at 4°C until use. Vibratome sections - Sample preparation Timing: 30 mins per sample
2. Fill sample tray of the vibratome with 0.15 M cacodylate buffer.
3. Prepare a collection well-plate or sample vials containing cacodylate buffer CAUTION-Cacodylate is toxic. Wear gloves at all times. CRITICAL STEP-If the free-floating slices will be immunolabeled and imaged with fluorescence the samples can be collected in TBS as this buffer has better spectral properties for imaging.
4. Put enough cyanoacrylate glue on the sample plate of the vibratome as needed to fully cover the base of the sample and attach it securely to the plate.
5. Place the brain block on the glue. Nudge the block with a paintbrush after a few seconds to ensure it is stable. CRITICAL-Sample should not dry out.
6. Immerse the sample plate into the sample tray containing the cacodylate buffer. The glue will harden upon immersion in the buffer.
7. Set amplitude and speed.
8. Set start and end points of the section.
9. Start cutting. CRITICAL STEP– We typically start with an amplitude of 0.8 and a speed of 0.6 mm/second. However, these parameters may need to be adjusted to find a speed and amplitude that lead to consistent cuts. Decreasing speed and increasing amplitude allows cutting of softer tissue and thinner sections. The desired thickness depends on the scientific question and technical aspects (such as size of ROI, antibody penetration, flat embedding etc).
10. Use a brush to collect sections into storage vials/well plate containing storage buffer.
11. Remove sample tray and decant the cacodylate buffer with tissue debris into a prepared waste container (remove big pieces of brain if necessary). CAUTION-Brain waste can be considered as bio-hazardous. Follow the internal safety rules of your facility/institute regarding waste disposal.
12. Remove the leftover glued brain by carefully sliding a razor blade between the sample and the support.
13. Peel off the attached glue from the tissue and discard.
14. Put the leftover tissue back in the storage buffer.
15. Store vibratome sections in the collection buffer until use at 4°C. Immunolabeling of free-floating sections Timing: 3 hours to overnight (depending on antibody incubation time) CRITICAL - Include appropriate controls to rule-out false positive signals and non-specific labeling (e.g. omitting primary antibody, empty channels to distinguish background/autofluorescence, isotype control). Fluorescence staining may require pre-testing of antibody concentrations and incubation parameters. Antigen retrieval could be performed here however the effect on the ultrastructure would need to be carefully assessed.
16. Wash the free-floating sections with three exchanges of 1x TBS for 5 minutes each at RT on orbital shaker.
17. Prepare the primary antibody cocktail in 1x TBS.
18. Immerse the sections in the primary antibody cocktail. Use enough primary antibody to adequately cover the sections.
19. Agitate the samples on an orbital shaker for the desired incubation time.
20. Wash the free-floating sections with three exchanges of 1x TBS for 5 minutes each at RT on orbital shaker.
21. Incubate the sections in secondary antibodies coupled to fluorescent dyes for 1h at RT on an orbital shaker.
22. Wash the free-floating sections with three exchanges of 1x TBS for 5 minutes each at RT on orbital shaker.
23. Store the sections in 1x TBS at 4°C for imaging. CRITICAL STEP-If samples are stored long term, it is best to add 0.1% final concentration of PFA to 1x TBS to preserve the ultrastructure (in analogy to the cacodylate storage buffer). Fluorescent imaging and ROI mapping Timing: Up to several hours per tissue section depending on the number of ROIs and size of area to be imaged. CRITICAL This section includes a general outline with the key points needed for creating a fluorescent map to be used for later correlation (Figure 2).
24. Reversibly mount the section on a glass slide.

a. Place a drop of mounting solution (50% Glycerol, 50% TBS v/v) onto a glass slide
b. Pick up a free-floating section using a brush and carefully roll the section off the brush and into the mounting medium on the glass slide
c. Manipulate the section with the brush to ensure it is completely flat and devoid of wrinkles or folds.
d. Place a precision coverglass directly on top of the section avoiding the creation of air bubbles over the sample.
e. Use a tissue to wick away excess liquid and secure the coverglass onto the glass slide.
25. Insert the glass slide into the microscope slide holder.
26. Take a low magnification overview of the entire section (e.g. using a 10x objective) in brightfield and fluorescence.
27. Image the ROIs at higher magnification if needed (e.g. for characterization purposes).
28. Adjust laser intensity, aperture and exposure to get an appropriate fluorescent signal for all channels at the magnification to be used for imaging.
29. Determine upper and lower z values of the ROI.
30. Record the z-stack.
31. Annotate the position of each imaged ROI as accurately as possible on the brightfield/fluorescent overview image. CRITICAL STEP-Some imaging softwares have integrated options to do this. Otherwise, the overview image can be opened in an external software and manually added.
32. Continue imaging and annotating ROIs as necessary.
33. Unmount the tissue section:

a. Immerse the glass slide in a 50 ml tube containing 1x TBS and wait for the coverslip to detach (∼5 mins). CRITICAL STEP: physically forcing the detachment of the coverglass is not recommended as the sections can easily tear.
b. Retrieve tissue section with brush.
34. Store the section in 1x TBS ready for resin embedding. PAUSE POINT-Sections can be stored in storage buffer at 4°C for a few weeks. *En bloc* staining and resin embedding Timing: 9.5 - 22.5 hours plus 48 hours for resin curing. CRITICAL This section details the full protocol for staining and resin infiltration and flat embedding of free-floating vibratome sections. Depending on the application (e.g. TEM only) a shorter protocol omitting some of the staining steps can be used. For SEM, the full protocol is recommended (Figure 3). For large samples (e.g. skin biopsies) the incubation times may need to be increased depending on the thickness of the sample. CAUTION-For the most consistent infiltration of solutions into the tissue, all steps should be performed with agitation unless otherwise stated. CAUTION-This step requires many hazardous substances therefore all steps should be performed in a fume hood. CRITICAL - Steps 37-39, 42-47 and 52-53 can be omitted for short protocol.
35. Wash the samples with multiple exchanges of cold 0.15 M CaCO buffer for a total of 15 minutes to remove all traces of chemical fixative (at least 3 exchanges of 5 minutes each).
36. Post-fix the samples by incubation in reduced osmium for 1 hour on ice. (Can be included in the short protocol for better preservation of membranes)
CRITICAL STEP-Prepare fresh TCH during this incubation step.
37. Wash the samples with multiple exchanges of ddH_2_O at room-temperature for a total of 15 minutes (e.g. at least 3 exchanges of 5 minutes each).
38. Immerse the samples in filtered TCH for 20 minutes.
39. Wash the samples with multiple exchanges of ddH_2_O at room-temperature for a total of 15 minutes (e.g. at least 3 exchanges of 5 minutes each).
40. Post-fix the samples by incubation in 2% Osmium tetroxide in ddH_2_O for 30 minutes at room temperature.
41. Wash the samples with multiple exchanges of ddH_2_O at room-temperature for a total of 15 minutes
42. Immerse the samples in 2% filtered uranyl acetate in ddH_2_O and leave for 1 hour at room temperature CRITICAL STEP-Uranyl acetate infiltration for 30 mins can be optionally included for increased contrast when using the short protocol. Prepare aspartic acid for the following day. PAUSE POINT-The samples can be left at 4°C overnight.
43. Wash the samples with multiple exchanges of ddH_2_O at room-temperature for a total of 15 minutes (e.g. at least 3 exchanges of 5 minutes each)
44. Bring samples to 60°C by incubation in a water bath or oven for 30 minutes.
45. Prepare Waltons lead aspartate at pH 5.5, 60°C and stabilize at 60°C for 30 minutes in a water bath.
46. Immerse the samples in Waltons lead aspartate and incubate at 60°C for 40 mins without agitation. CRITICAL STEP-Prepare the Durcupan resin during this step.
47. Wash the samples with multiple exchanges of ddH_2_O at room-temperature for a total of 15 minutes (e.g. at least 3 exchanges of 5 minutes each).
48. Dehydrate the samples using a graded alcohol series (20%, 50%, 70%, 90%, 3 x 100%) through immersion for 5 minutes each at room-temperature. CRITICAL STEP-Samples will become brittle after alcohol dehydration so care should be taken when handling them after this point.
49. Immerse the samples in 50% resin in alcohol for 30 minutes at RT.
50. Exchange the samples into 100% resin for 1 hour.
51. Exchange the samples into fresh 100% resin and leave to infiltrate overnight. CRITICAL STEP-It is critical that the samples are agitating well (but not to cause air-bubbles in the resin or cause the brittle samples to break) so they do not stick to the bottom of the vials/well plate during this long incubation period. Infiltration time can be reduced to 4 hours when using the short protocol.
52. Prepare fresh Durcupan resin.
53. Exchange the samples into 100% resin and incubate for 2 hours. CRITICAL STEP-Use the excess resin to fill flat embedding molds to make resin stubs for mounting samples in preparation for ultramicrotomy.
54. Flat embed the samples

i. Prepare a glass-slide and Aclar sandwich. CRITICAL STEP: Make sure the Aclar sheets are bigger in all dimensions than the glass slide to avoid the glass slides being stuck together by the cured resin.
ii. Place a drop of resin on one of the Aclar pieces.
iii. Use a toothpick or cut a diagonal surface on the end of an applicator stick to pick up and manipulate the samples in the resin.
iv. Pick up the slices and drop onto the Aclar film being careful not to break the samples.
v. Add some additional resin surrounding the sample. CRITICAL STEP-Don’t add so much resin that it bleeds out over the edges of the Aclar when the sandwich is placed on top, but enough so that the Aclar sticks.
vi. Roll the second piece of Aclar film carefully against the tissue slices in a way that avoids air bubble formation.
vii. Sandwich the Aclar between two glass slides.
55. Cure the resin in a 60°C oven for 48 hours. Tissue block preparation Timing: 1-2 hours CRITICAL The regions in the sample containing the ROIs must be excised out of the sample and mounted on a resin stub in preparation for serial sectioning (Figure 4).
56. Take a brightfield image of the entire resin-embedded section.
57. Overlay the image of the resin section with the fluoro/brightfield map in any suitable software (e.g. ppt/illustrator etc.). CRITICAL STEP-Ensure the scale is set accurately.
58. Draw trapezoid shapes around the ROIs with an area that will maximally fit on a 1 x 2 mm slot grid (e.g. – 1.5 x 0.7 mm) CRITICAL STEP-The trapezoid shape helps for sample orientation in the downstream EM imaging. Try to put ROIs in the center of the shape away from the edge.
59. Draw and overlay a numbered grid on top of the shape design.
60. Peel the top Aclar sheet off the embedded resin sections.
61. Print the grid and shape design (to scale) onto transparent paper and overlay onto the embedded section CRITICAL STEP: Using the edges of the section and other physical features of the tissue will help with the alignment.
62. Under binoculars, use a razor blade/scalpel to excise the tissue blocks. (Optional) Excise shapes using laser cutting (Supplementary Figure 5)
63. Remove excised tissue blocks by placing sticky tape over the section and slowly peeling it back. The cut shapes should adhere easily to the tape.
64. Mount excised tissue blocks on a resin block (steps 65-67). CRITICAL STEP: a rectangular shape may be easier to excise from the tissue. In this case, use a razor blade/scalpel to trim the block surface area into a trapezoid shape after it is mounted on the resin block.
65. Spread super glue on top of an empty resin block. CRITICAL STEP-The top surface of the empty resin block can first be scratched with sandpaper to create a flat surface with better adherence for the tissue. CAUTION-Resin particles created by sandpapering can be a skin irritant. Perform this step in a fume hood and wear protective clothing.
66. Pick up the excised block with a wetted toothpick and mount on top of the prepared empty resin block.
67. Wick away the excess glue taking care not to dislodge the sample. CRITICAL STEP-The glue must dry completely to ensure it is stable for ultramicrotomy. This step can be done the night before or a few hours prior to the cutting session. Serial sectioning and collection Timing: 2 hours per sample
68. Place the resin block on the sample holder and use a razor blade to trim away empty resin on the support resin block.
69. Place the sample on the ultramicrotome arm sample holder
70. Mount the diamond knife on the ultramicrotome (Figure 5A). CRITICAL STEP-If there is too much excess resin on top of the sample, this can be first trimmed away using a trimming knife. The trimmed resin will look white on the knife. As soon as the knife reaches the sample in the resin block, the trimmed parts look slightly darker. However, caution should be used to ensure not too much sample is trimmed away in case the ROI is located at the top of the block in z-height.
71. Align the knife edge to the block surface. CRITICAL STEP-The correct alignment of the block to the knife is necessary to minimize sample and region of interest loss during the cutting.
72. Approach the diamond knife to the sample until the light reflection on the sample surface is barely seen.
73. Set the following parameters on the ultramicrotome:

a. Cutting window (start/end cutting point)
b. Sample thickness (variable (50-200 nm)
c. Cutting speed (1.6 mm/s)
d. Activate the counter mode to have an estimation of how much resin is cut
74. Overfill the boat of the diamond knife with filtered ddH2O to make sure the entire surface of the knife edge is wet.
75. Use a small syringe fitted with a hypodermic needle to remove excess water until the water surface becomes shiny and flat like a mirror. CRITICAL STEP-The diamond knife must always stay hydrated while cutting, otherwise the sections will shrink and fold.
76. Start cutting some sections to check if all the empty resin has been removed. CRITICAL STEP-Usually the resin-only section is very smooth and uniform while if it contains tissue some pattern is visible on the surface of the section, and it can appear rough.
77. (*Optional*) To check whether the section contains any sample or if it is resin-only,

a. collect 1-2 sections on a glass slide and stain with toluidine blue:

i. Use the eyelash manipulators to isolate the section (s) of interest.
ii. Use the loop to pick up 1-2 sections.
iii. Place them on a clean glass slide.
iv. Let the water droplet dry by placing the glass slide on the hot plate.
v. Cover the section (s) with a droplet of toluidine blue solution.
vi. Heat the slide on the hot plate (at roughly 100°C) for about 30 seconds, or until the droplet edge dries (turns green).
vii. Rinse with ddH2O until the droplet edge has been cleared.
viii. Dry the slide on the hot plate for a few seconds until the water evaporates.
ix. Inspect the slide with a light microscope. CRITICAL STEP-If the section contains any sample, cellular and extracellular components will now be visible, otherwise the section will remain transparent (Figure 5B).
78. If necessary, repeat steps 77 until the sample or the region of interest becomes visible in the section.
79. When ready for collection prepare the following material:

a. Use the diamond scribing tool to scratch the bottom surface of some glass slides to create reference positions, to later unambiguously identify the sections in the correct order (Figure 5C).
b. Prepare some tweezers with EM grids according to the collection strategy. CRITICAL STEP-The order of the sections during the collection might be very important so make sure to have all the tools ready and prepared for the serial collection.
80. Cut a ribbon containing 14 sections for one CLEM cycle as follows: 6 semi-thin sections (200 nm) for glass slides, 7 ultra-thin sections (<100 nm) for EM grids, 1 semi-thin section (200 nm) to be left on the diamond knife for the next cycle. CRITICAL STEP-Sections of different thicknesses have different colours (blue/green - 180-250 nm, purple 150 – 170 nm, gold - 90-140 nm, silver - 60-80 nm) (Figure 5A). This is helpful to remember which sections need to be placed on glass slides and on EM grids. The collection as described here will span approximately 1.5 µm depth in 1 CLEM cycle. For collection all the way through a 20 µm ROI/cell, 18 CLEM cycles should be collected. The number and thickness of the collected sections can be changed according to the end-requirements. E.g. - thicker EM sections (120-200 nm) could be collected for tomography and less, or thinner, sections in the CLEM cycle for a smaller ROI. CRITICAL STEP-Consider collecting multiple glass slides in parallel, to have multiple IHC labeling possibilities and/or to include toluidine blue staining on one slide
81. Transfer the sections on glass slides/grids in the correct collection order (Figure 5C).

a. Use the eyelash manipulators to detach the ribbon from the knife leaving the last-cut semi-thin section on the knife, separate the section (s) of interest into groups as they will be collected on the glass slides and EM grids.
b. Use the loop to pick up the first 3 sections and place them on the first glass slide in position 1.
c. Use filter paper to wick away the water from the loop leaving the sections flat on the glass slide.
d. Use the loop to pick up the second 3 sections and place them on the second glass slide in position 1.
e. Immerse an EM grid in the water bath and position it underneath a group of ultra-thin sections (for a mesh grid) or a single isolated section (for a slot grid)
f. Use the eyelash manipulators to slide the sections on the grids while simultaneously lifting the grid out of the water bath. CRITICAL STEP-Sections (especially the thin ones) are very prone to breakage. Hold the tweezer with the grid with one hand and keep it underwater near the selected section and with the other hand use the eyelash manipulator to gently move and place it on the grid. Once the section is on the grid, move the tweezers out of the boat and let the water droplets on the grid dry.
82. Once the collection is finished, place the glass slide on the hot plate for 15 mins so that all the water dries out and the sections properly adhere to the glass slide.
83. Store all the grids in an empty grid box. PAUSE POINT-The sections mounted on both the glass slides and EM grids can be stored indefinitely at room-temperature until needed. Toluidine Blue staining Timing: 1-12 hours CRITICAL Counterstaining a glass slide with toluidine blue can help with the tissue morphology identification and for sample correlation/alignment purposes. (Figure 6A)
84. Cover the sample on the glass slide with a droplet of toluidine blue solution.
85. Heat the slide on the hot plate (at roughly 100°C) for about 30 seconds, or until the droplet edge dries (turns green).
86. Rinse with ddH2O until the droplet edge has been cleared.
87. Dry the slide on the hot plate for a few seconds until the water evaporates.
88. Use a mounting medium to coverslip the slides.
89. Leave the slides in the oven for 1 hour at 37°C or overnight at RT to let the mounting medium dry. Immunohistochemistry (Optional) Timing: 1 day CRITICAL-Through the whole protocol don’t let the slides dry. If many slides are stained at the same time, process them in batches. Be careful not to scratch the surface of the section during the manual handling of the slides throughout this section. Include appropriate controls to rule-out false positive signals and non-specific labeling (e.g. omitting primary antibody, isotype control).
90. Prepare the etching solution in a glass staining jar at least 12 hours before the etching. For the etching, place the glass slides into the glass staining jar containing the etching solution and etch for 3 minutes. CRITICAL STEP-Use a dedicated waste for disposal of the etching solution.
91. Rinse glass slides in 100% Ethanol. Pre-treatment/antigen retrieval Timing: 45 minutes
92. Rinse slides in ddH2O.
93. Put glass slides in 100% Formic acid for 10 min. CRITICAL STEP-Work under the fume-hood.
94. Rinse slides in ddH2O.
95. Put glass slides in 1 x Tris-EDTA, pH 9 or 1 x NA-Citrate, pH 6 (depending on the antibody) for 30 minutes at 100°C. CRITICAL STEP-Use a plastic staining jar with a lid to avoid evaporation and place it in the steamer.
96. Wash slides in ddH2O to cool down the slides. Blocking Timing: 30 minutes
97. Put slides in the endogenous HRP blocking solution for 10 minutes at RT.
98. Wash slides in PBS 3 times (quick wash, quick wash, 5 minutes) PAUSE POINT-Slides can stay in PBS for a few days.
99. Place all the slides on the staining tray and use a pathology pen to draw an area around the sections on the glass slide to contain the staining. CRITICAL STEP-Put some water in the bottom of the staining tray to keep the environment moist.
100. Add a few drops of antibody diluent inside the area and leave it for 10 minutes. Antibody staining Timing: 2-12 hours
101. Prepare the correct dilution of the primary antibody in the antibody diluent. CRITICAL STEP-Primary antibody concentration, timing, incubation temperature, and antigen retrieval are highly dependent on the specific antibody and needs to be optimized prior. Common time and temperature combinations are 1 hour at 37°C, 4 hours at RT, overnight at 4°C.
102. Incubate the primary antibody according to its working time/temperature combination. CRITICAL STEP-Prepare PBS-T solution during primary antibody incubation.
103. Wash slides 3 times in PBS-T (quick rinse, quick rinse, 5 minutes incubation)
104. Dry the pathology pen marks on the glass slides by rolling the end of a cotton tip over the surface of the pen mark. NOTE-The detergent stops the pathology pen creating a liquid boundary. Avoid touching the area where the sections are.
105. Add a few drops of secondary antibody on glass slides and incubate for 30 minutes at RT. CRITICAL STEP-Make sure to match primary and secondary antibodies species.
106. Wash slides 3 times in PBS (quick rinse, quick rinse, 5 minutes incubation). PAUSE POINT-Slides can stay in PBS for a few days.
107. Prepare HRP-chromogen detection solution.
108. Stain slides with HRP-chromogen solution for 3 minutes at RT.
109. Rinse slides in 1 x PBS.
110. Place slides in ddH2O to stop the reaction.
111. Contrast the slides with hematoxylin staining (protocol in Supplementary Information)
112. Repeat steps 88 and 89 to coverslip the slides. PAUSE POINT-The slides can be dried at this point by inverting and gently pressing against some absorbent material. Dried slides can be kept for a few weeks without any changes. Imaging of stained glass slides Timing: 2 hours per sample
113. Image the glass slides processed for IHC and toluidine blue with a light microscope at the magnification necessary to see the tissue features and location of stained ROI.
114. Annotate cycles containing ROIs showing positive IHC staining. They will be used to find the corresponding EM grids in the next step.
115. The IHC and toluidine blue images for each cycle can be aligned here using FIJI BigWarp to follow the ROI through the block (steps 116-125). Image correlation using BigWarp Timing: 5 minutes CRITICAL In this section we perform a simple alignment between two images (Figure 7A).
116. Open FIJI.
117. Load the images to be aligned CRITICAL STEP-A rough alignment can first be performed by flipping and/or rotating the images. To do so, click Image >Transform.
118. Click Plugins > BigDataViewer > Big Warp.
119. Define which image will be the moving and target image. For example:

a. IHC (moving) onto toluidine blue (target)
b. IF (moving) onto toluidine blue (target)
120. Press the spacebar to toggle the landmark mode on/off.
121. Click a point on one image and click the same point on the other image to add the landmarks. Add at least 4 landmarks.
122. (Optional) File > save landmarks.
123. Press T to transform the moving image.
124. Click File > Export moving Image.
125. Save the resulting image. CRITICAL STEP-Check the image alignment by making the images into stacks. Click Image > Stack > Images to Stack.

**Figure 2.**
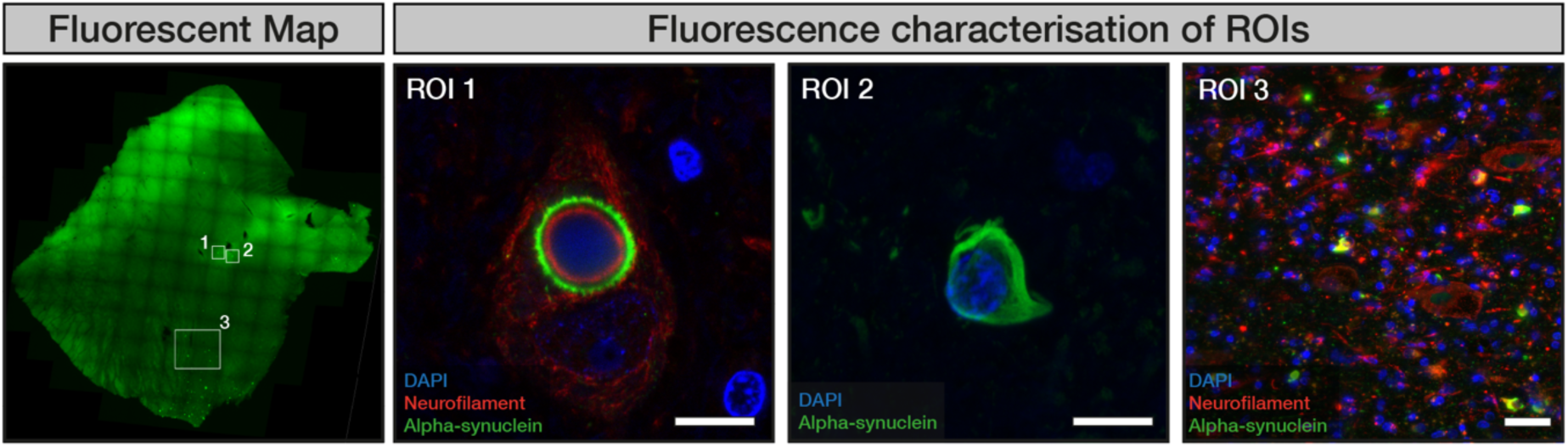
Fluorescent mapping of free-floating sections. A fluorescent overview of an entire brain slice is taken at low magnification. Individual cells or areas (white square) to be targeted for EM are annotated as accurately as possible on the overview image. High resolution images and/or volumetric z-stacks of the annotated ROIs can be obtained. ROIs 1-3 depict alpha-synuclein disease pathology in order: a single neuron containing a Lewy body, the pathological hallmark of Parkinson’s disease; a single glial cytoplasmic inclusion, the pathological hallmark of multiple systems atrophy; a large region showing multiple alpha-synuclein inclusions alongside neurons from the brain of a multiple systems atrophy patient. For each image, the maximum fluorescence projections of volumetric z-stacks are shown, deconvoluted using the THUNDER algorithm (Leica). DAPI staining for cell nuclei plus antibody-specific immunostaining as stated are shown. Scale bar is 10 µm for ROI 1 and ROI 2, and 20 µm for ROI 3. These images were produced during the data collection from^10^. Images adjusted for contrast.

**Figure 3.**
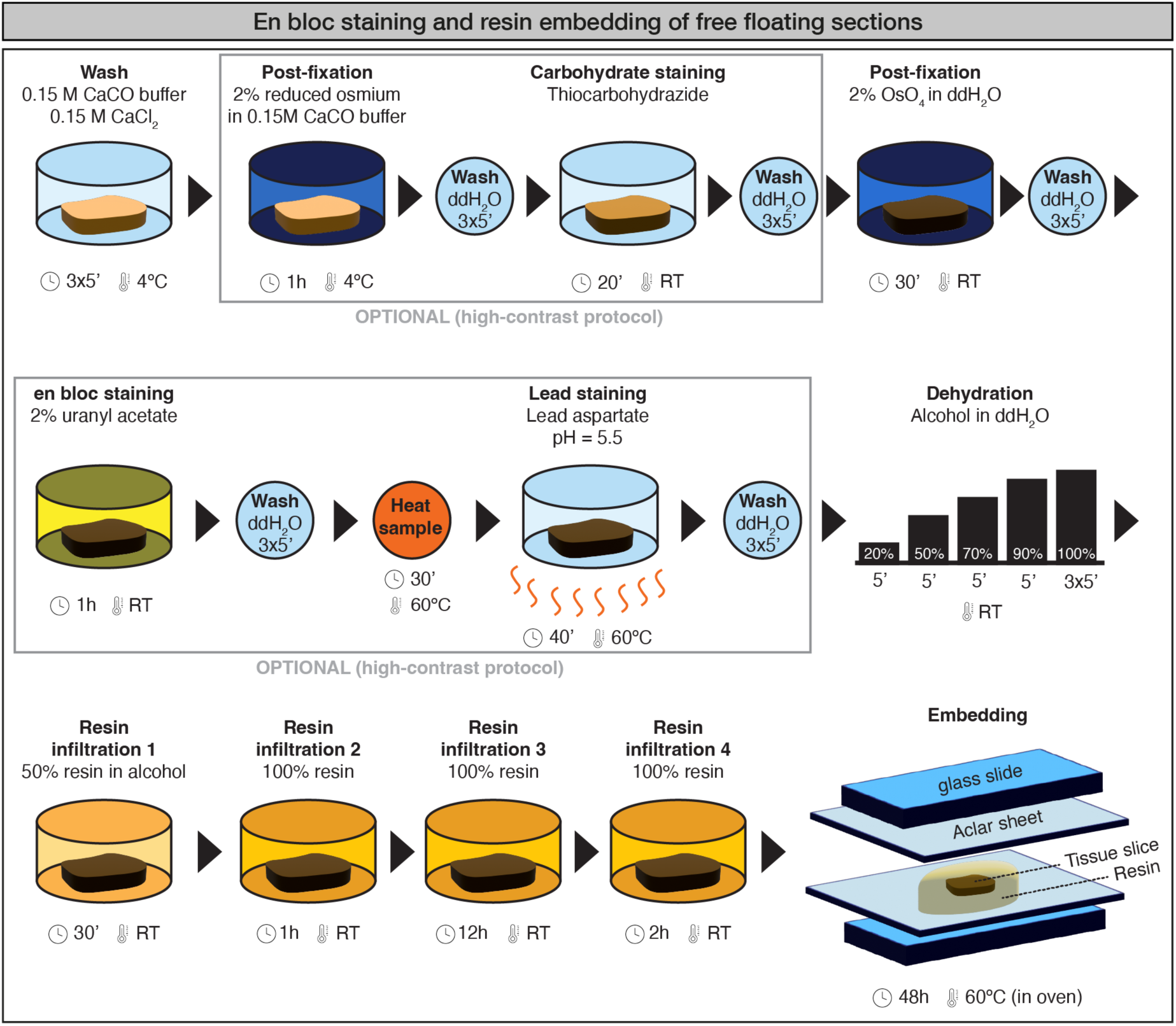
Schematic of resin embedding process on free-floating sections.

**Figure 4.**
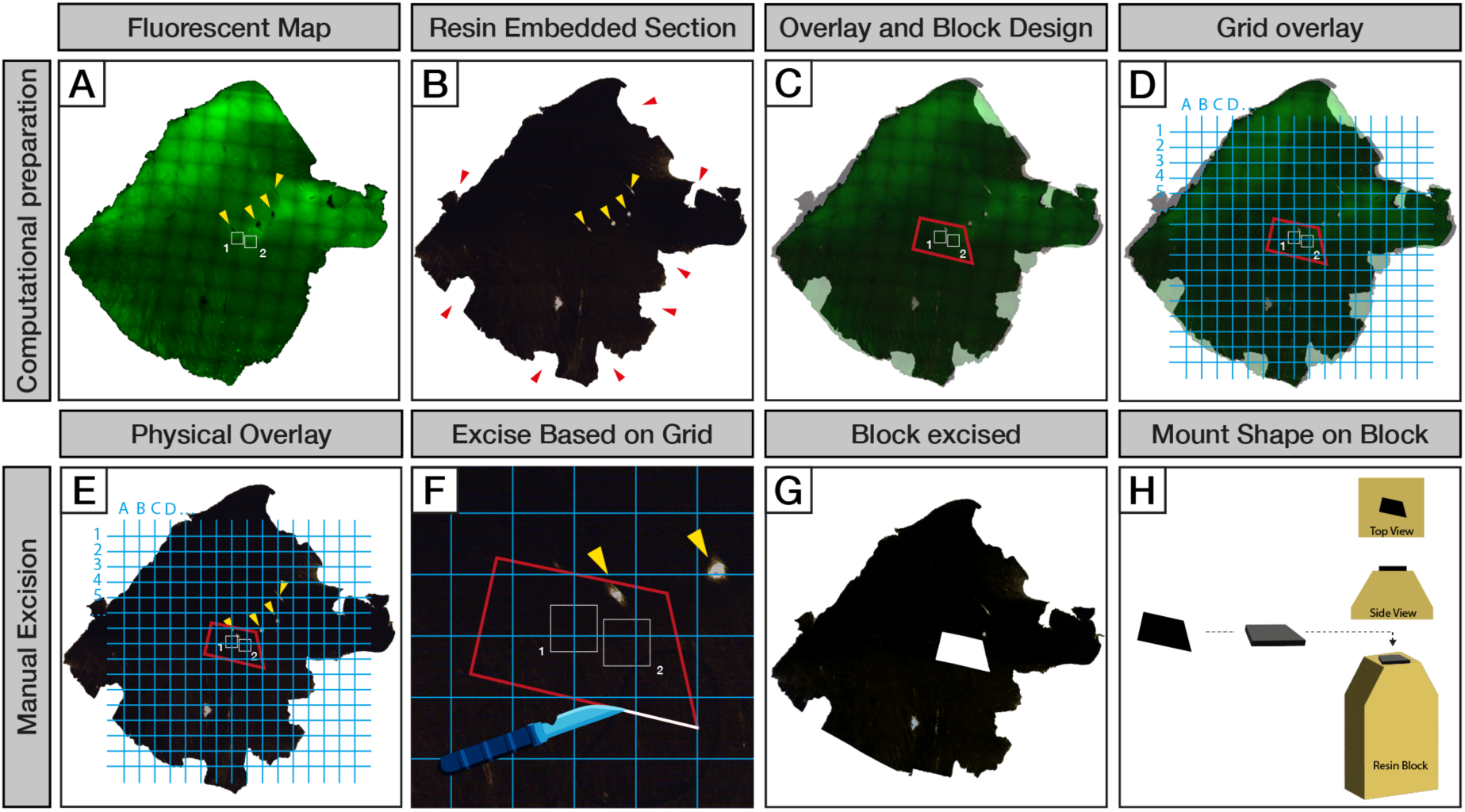
Preparation of blocks for CLEM sectioning. The fluorescent map made from the free-floating section (A) is shown next to a brightfield image of the resin embedded section (B). The edges of the resin section have been broken during the embedding process (red arrows) however, macro features visible in both images (yellow arrows) can be used for a precise alignment around the position of the ROIs (white squares). The two images are overlaid computationally, and a trapezoid shape (red outline) is drawn around the ROIs (C). A grid is overlaid onto the ROI map to identify the coordinates needed for the shape excision (D). The grid and ROI map are printed to scale and physically overlaid with the resin embedded section (E). The trapezoid tissue block is excised from the resin-embedded section using a scalpel (F, G), and mounted onto a resin block (H). Images were taken on a THUNDER Imager 3D Tissue (Leica) equipped with a K8 fluorescent camera (fluorescence) or a DMI8 colour camera (brightfield). The background was digitally subtracted from the raw images for better alignment of the section edges.

**Figure 5.**
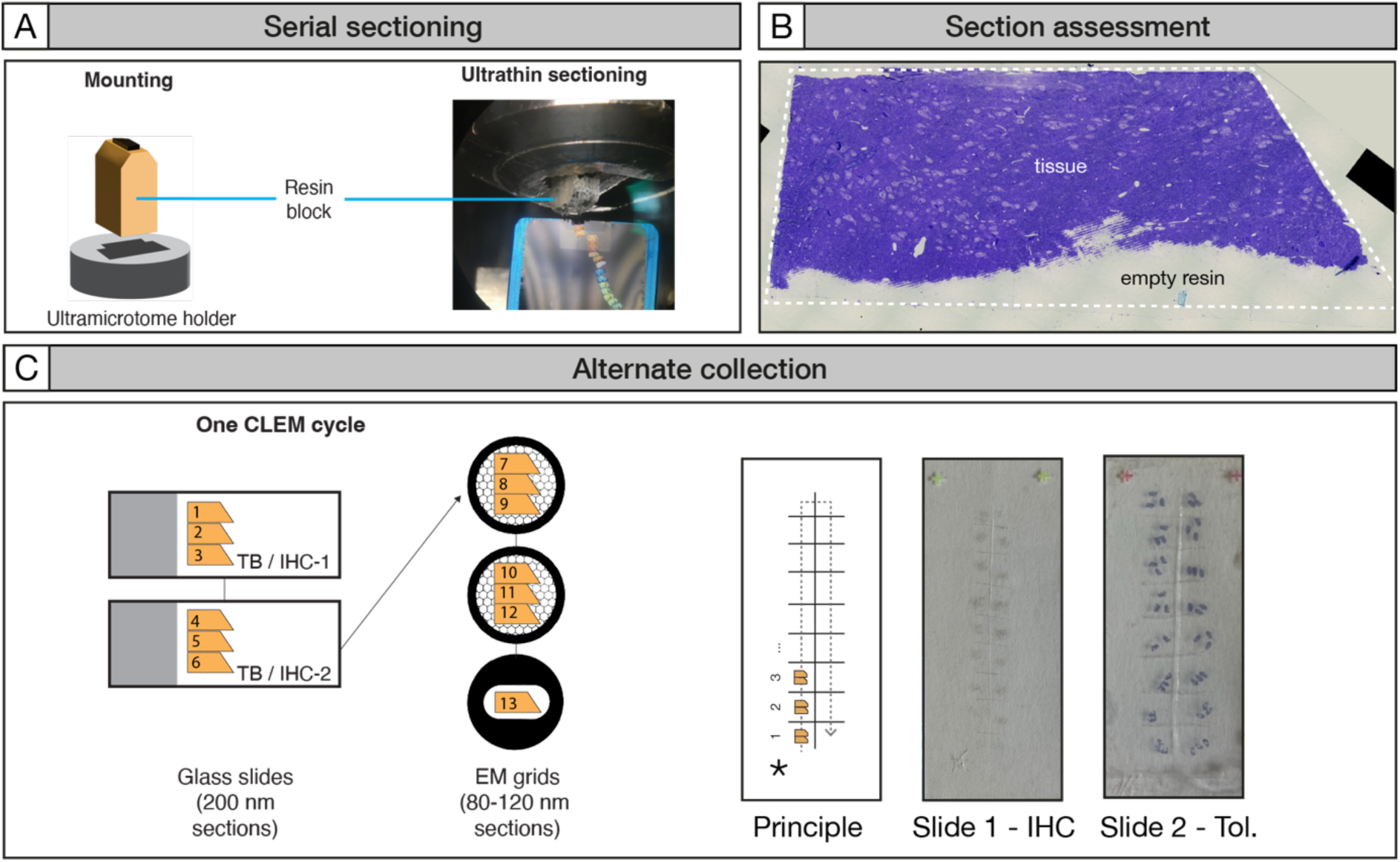
CLEM sectioning and IHC staining of brain tissue sections on glass slides. The prepared resin blocks are mounted onto an ultramicrotome. Serial ultra-thin sections (80-200 nm) are collected from the blocks. Sections of different thicknesses will refract different colours (A). Toluidine blue staining can be performed on-the-fly to assess the proportion of tissue/resin on the cut sections (B). The sections are collected alternatively on EM grids and glass slides in cycles all the way through the sample. Sections within each CLEM cycle can be collected sequentially on multiple glass slides to be used for toluidine blue staining and/or multiple antibodies in IHC (C-left). Schematic showing the markings on the underside of the glass slide etched with a diamond scribe (C-right). The grid delineates where the sections should be placed for each collection cycle and a mark (*) denoting the start position. The numbers indicate the order of each CLEM cycle collected. Example images of unstained sections (Slide 1-IHC) and toluidine blue stained sections (Slide 2-Tol.) placed on the glass slides are shown.

**Figure 6.**
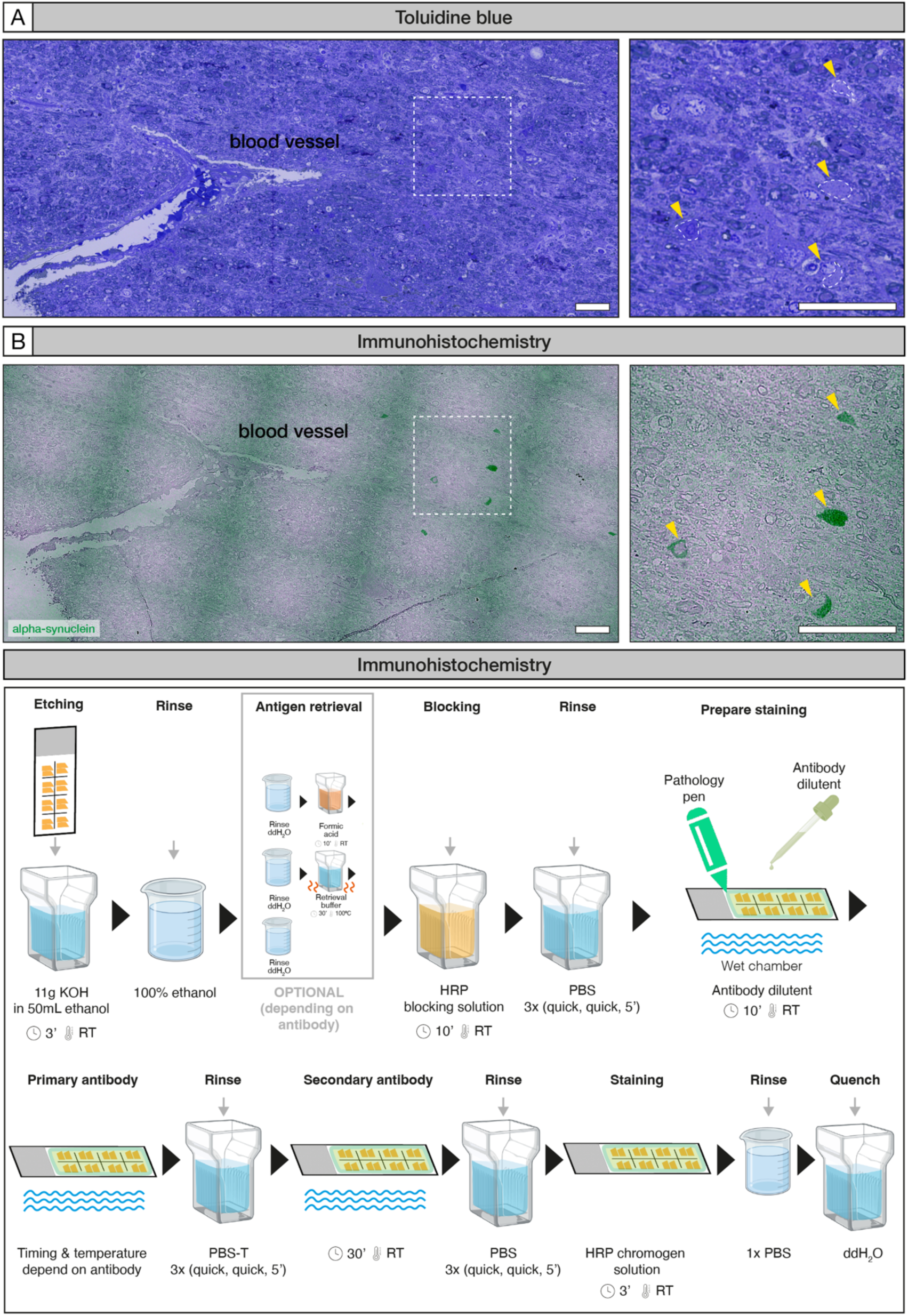
Staining of glass slides. The glass slides can be stained with toluidine blue (A) and/or IHC (B) to find which cycle contains the ROI (yellow arrows). The same ROIs identified in adjacent ultra-thin sections stained with either Toluidine blue (white outline) or IHC (green) are shown. These images were produced during the data collection from^11^. IHC image is colour adjusted and the contrast strongly increased to increase visibility. Scale bars are 1 µm. Schematic summary outlining IHC staining protocol of glass slides (B).

**Figure 7.**
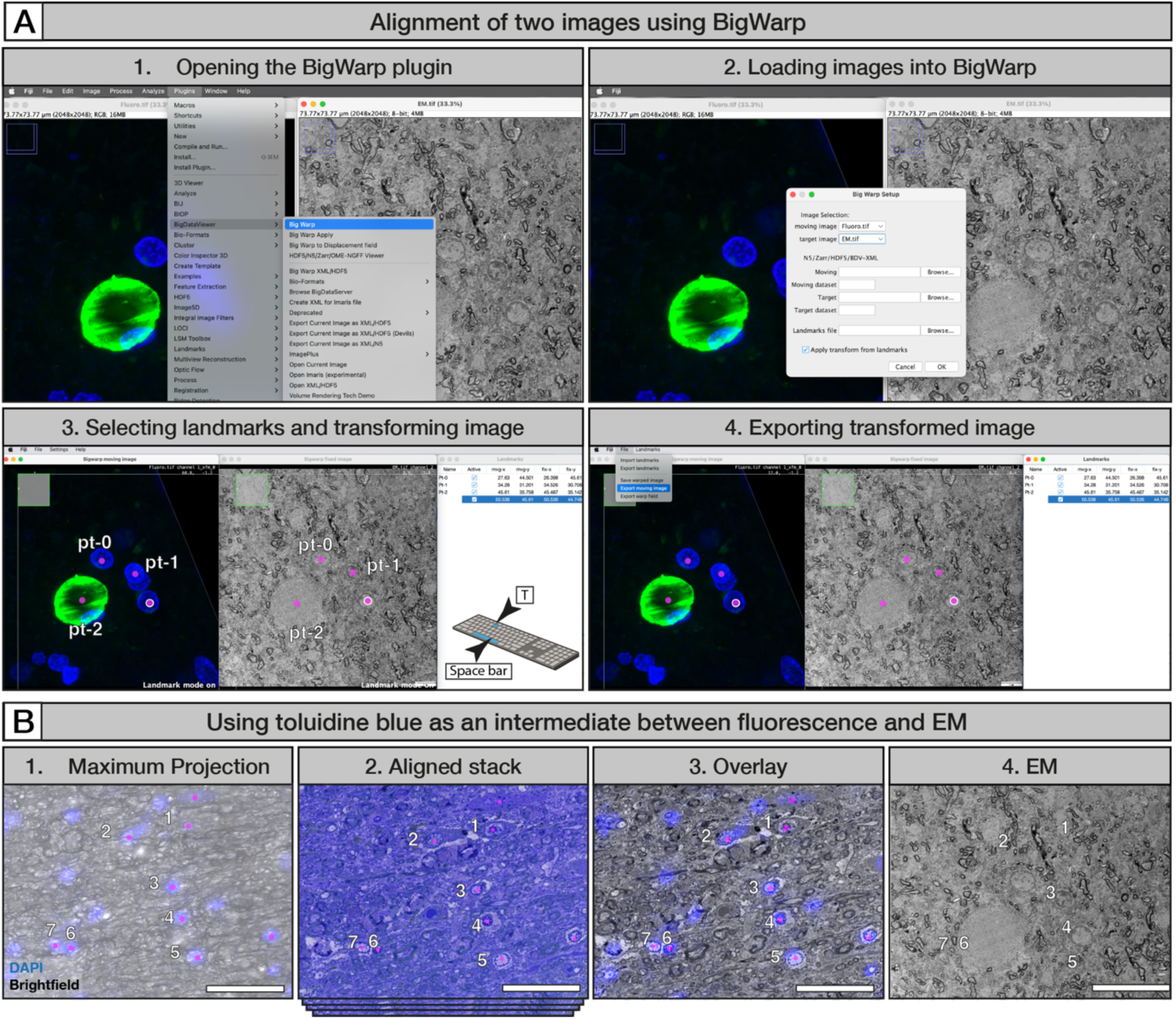
Image correlation using Big Warp. (A) Screenshots of a simple alignment of two images. 1) Two images are loaded into FIJI and the Big Warp plugin is opened. 2) The moving and target images are defined. 3) The space bar on the keyboard is pressed to toggle the landmark mode on and off. With the landmark mode on, identical features in on both images are selected. The coordinates of the selected landmarks are shown in a pop-up window. T is pressed on the keyboard to transform the moving image in relation to the target image. 4) The moving image is exported. These images were produced during the data collection from^10^. (B) Aligning a fluorescence volume with EM. 1) Fluorescence maximum projection of a 40 µm z-stack showing brightfield and DAPI channels (blue signal); 2) After resin embedding and serial sectioning, the brightfield images of the 200 nm toluidine blue stained sections on the glass slides are aligned into a z-stack; 3) Overlay of fluorescence maximum projection and toluidine blue stack (converted to black and white image) based on nucleus signal; 4) The electron micrograph of a grid from the same cycle shown in 1-3. The fluorescent maximum projection in (1) has been warped onto the toluidine blue image in (2) to identify exactly which grid should be imaged by EM. Pink dots indicate cell nuclei in the fluorescence volume that could be correlated on the toluidine blue section shown in (2). The numbers indicate a nucleus pattern that is visible in fluorescence, toluidine blue images, and electron microscopy. These images were produced during the data collection from^11^. Scale bar is 50 µm.

Fluorescent volume correlation using BigWarp Timing: 2 hours CRITICAL Correlation of a fluorescence volume taken on a free-floating section presents a challenge when aligning with an ultrathin 200 nm LM image or 80 nm TEM images. The fluorescent and LM/TEM images are often deformed in respect to each other in a non-linear way and some additional correlation steps are needed (Figure 7B).

1. Make a maximum projection of the fluorescence volume showing the DAPI and Brightfield channel. CRITICAL STEP-The brightfield channel is useful to provide tissue context to help orient the fluorescence sample with the toluidine blue/ IHC /EM images. Other channels/physical fiducial markers which would help with the orientation can be used instead/additionally.
2. Take a snapshot of the image with high enough resolution to resolve individual cell nuclei.
3. Align all the toluidine blue/IHC images acquired from all the CLEM cycles collected from the block volume (steps 77).

a. Align the image from cycle 1 (target) to the image from cycle 2 (moving).
b. Transform and save the moving image (cycle 2 transform).
c. Align the cycle 2 transform (target) to the image from cycle 3 (moving).
d. Transform and save the moving image (cycle 3 transform).
e. Continue until all the images have been transformed onto the image from the previous cycle.
4. Close all non-transformed images (except for cycle 1).
5. Click Image > Stack > Images to Stack.
6. Scroll through the stack to check the alignment and visualize the features as they move through the volume.
7. Align the fluorescence to the toluidine blue stack.

a. Separately open one image in the stack to serve as the toluidine reference image.
b. Open the fluorescence maximum projection.
c. Perform a basic alignment of the fluorescence (moving) and toluidine blue (target) using any features in the tissue that can be aligned confidently. (steps 114-123).
d. Press T to transform the fluorescent image.
e. Save the fluorescent transform.
f. Add the fluorescence transform to the aligned toluidine blue stack adjacent to the position of the reference toluidine blue image.
g. Move back and forth between the images to identify nuclei or other tissue features that correlate between the fluorescent transform and the reference toluidine blue images.
h. Scroll up and down the stack to identify features or cell nuclei that become visible in different planes within the stack.
i. Add the newly correlated nuclei/features as additional landmarks in BigWarp.
j. Repeat steps 130 d-i until the region surrounding the ROI has been correlated with a high level of confidence.
k. Save the landmarks.

CRITICAL STEP-The same transformation can be applied to different images. For example, if you first warp the fluorescence to toluidine blue (T1) and toluidine blue to EM (T2), you can directly warp the fluorescence to the EM by doing a T1>T2 transformation. In that case, you can only re-use the T2 transformation matrix if the pixel size of all images are set correctly, and the coordinates of the landmarks are saved in a unit that makes sense (so µm and not pixels). The pixel size can be adjusted in FIJI through the menu: Image > Properties. Use the same units for all images (e.g., μm).

EM image acquisition Timing: 1 hour

1. Find the EM grid corresponding to the cycle containing the positive IHC staining on the glass slide or recognized in the toluidine blue staining.
2. Load the EM grid in a TEM.
3. Use easy recognizable features in the tissue (e.g. cell nuclei, blood vessels, resin section edges) to locate the ROI (s) on the grid corresponding to the positive IHC signal (Figure 8).
4. (Optional) CLEM4CINA can be used to help locate the ROI on the grid (Supplementary Information; repository link https://github.com/LBEM-CH/clem4cina).
5. Acquire EM images / tomography as desired.
6. Correlate with FIJI BigWarp again as needed.

**Figure 8.**
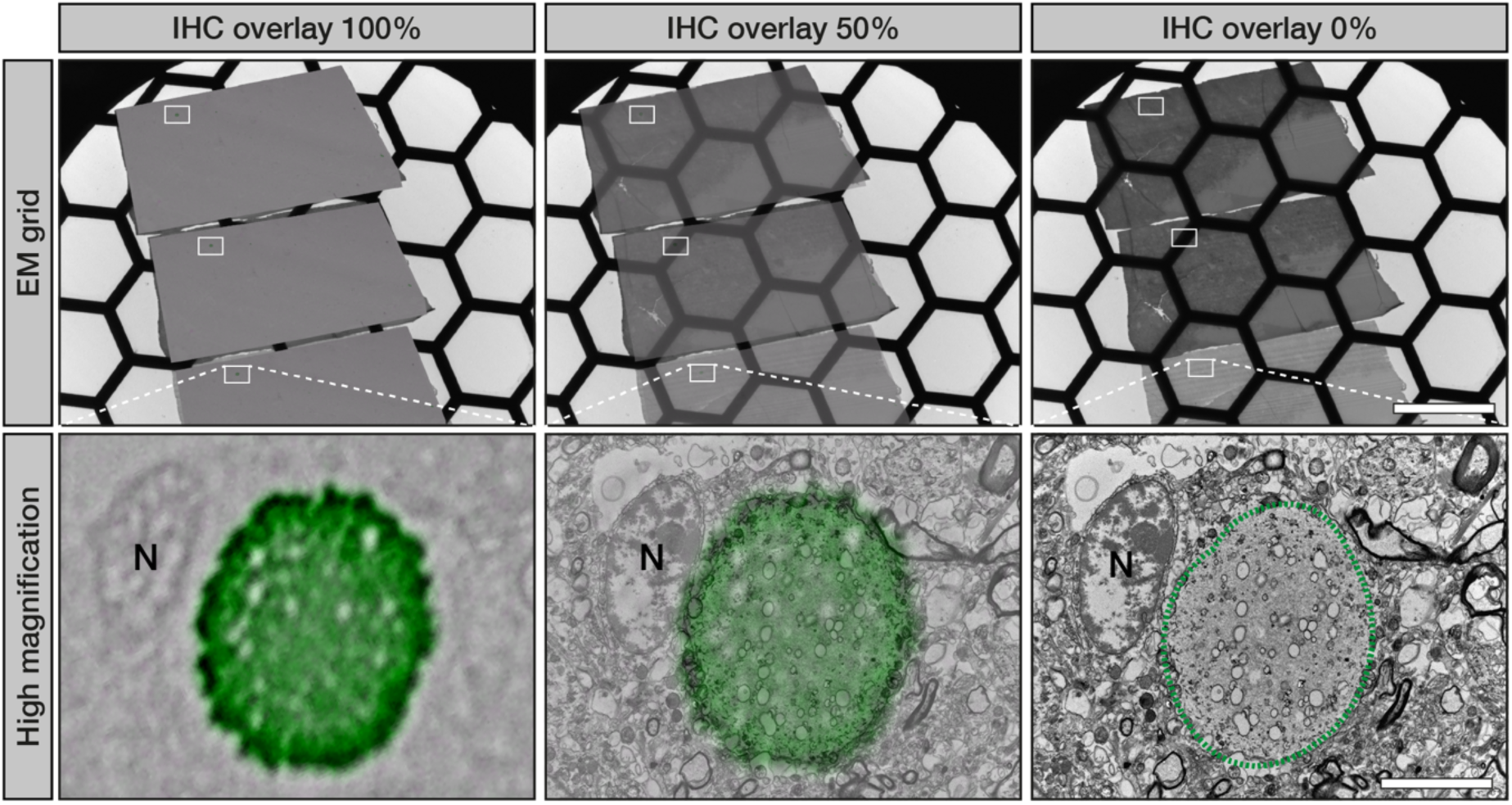
IHC and EM correlation. An ROI is located on the EM grid by overlaying a low magnification immunopositive brightfield IHC image with an overview of the serial sections on the EM grid within the same cycle. At higher magnifications, features visible in both imaging modalities, such as cell nuclei (N), can be used for exact immunopositive (green) ROI correlation. EM images were taken at room-temperature on a Tecnai G2 Spirit (ThermoFisher) operated at 80kV and equipped with a EMSIS Veleta camera. These images were produced during the data collection from^10^. Scale bar is 250 µm for EM grid overviews, and 5 µm for high magnification images.

#### Timing

In total approximately 7 days for pure experimental time needs to be accounted for. The total time spent will likely exceed this, as the timing is given for back-to-back experimental workflows. The timing does not include the reagent preparation.

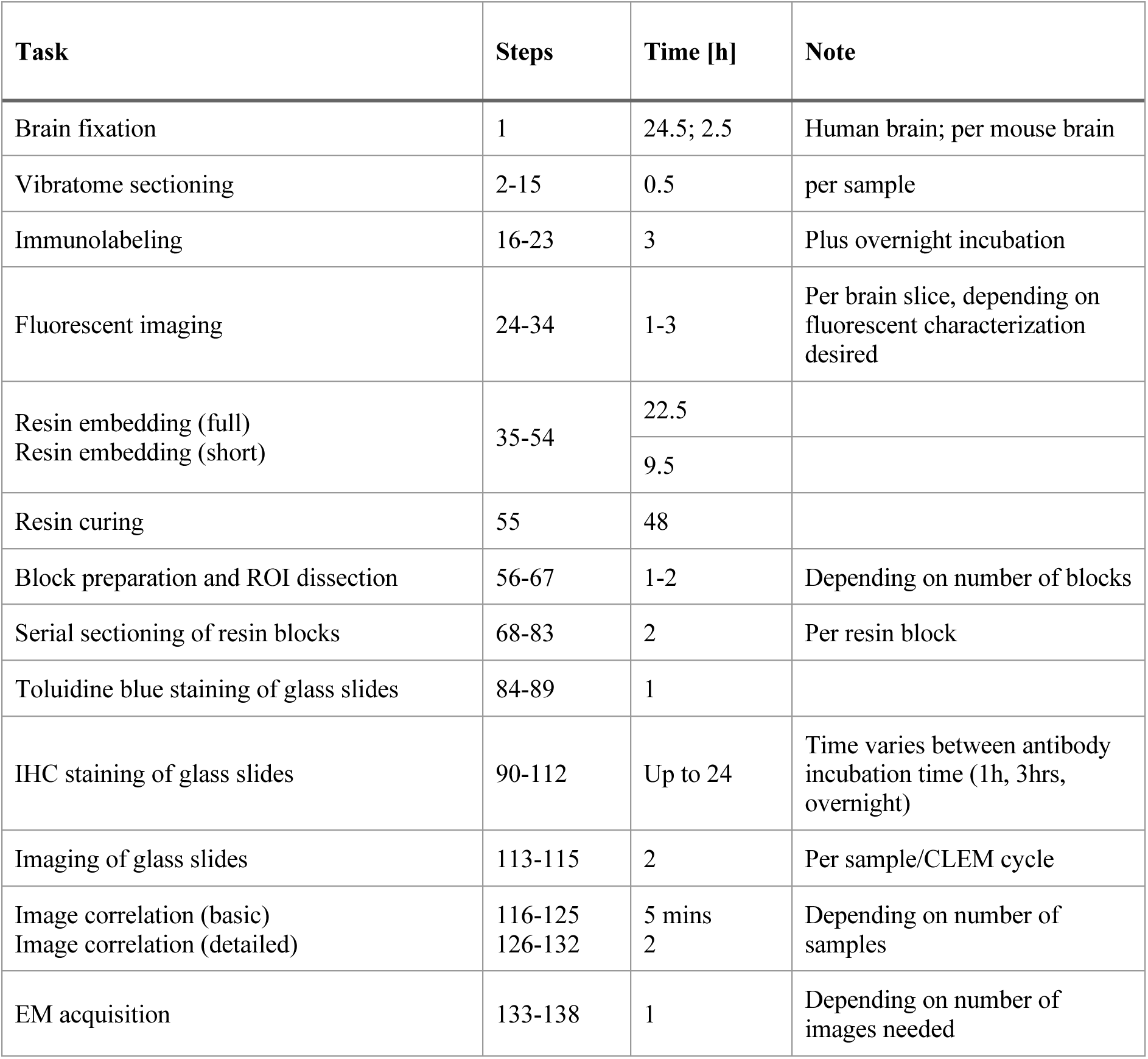

| Task | Steps | Time [h] | Note |
| --- | --- | --- | --- |
| Brain fixation | 1 | 24.5; 2.5 | Human brain; per mouse brain |
| Vibratome sectioning | 2-15 | 0.5 | per sample |
| Immunolabeling | 16-23 | 3 | Plus overnight incubation |
| Fluorescent imaging | 24-34 | 1-3 | Per brain slice, depending on fluorescent characterization desired |
| Resin embedding (full)<br>Resin embedding (short) | 35-54 | 22.5 |  |
|  |  | 9.5 |  |
| Resin curing | 55 | 48 |  |
| Block preparation and ROI dissection | 56-67 | 1-2 | Depending on number of blocks |
| Serial sectioning of resin blocks | 68-83 | 2 | Per resin block |
| Toluidine blue staining of glass slides | 84-89 | 1 |  |
| IHC staining of glass slides | 90-112 | Up to 24 | Time varies between antibody incubation time (1h, 3hrs, overnight) |
| Imaging of glass slides | 113-115 | 2 | Per sample/CLEM cycle |
| Image correlation (basic)<br>Image correlation (detailed) | 116-125 | 5 mins | Depending on number of samples |
|  | 126-132 | 2 |  |
| EM acquisition | 133-138 | 1 | Depending on number of images needed |

### Troubleshooting

| Step | Problem | Possible Reason | Solution |
| --- | --- | --- | --- |
| 26 | Fluorescence signal and resolution are too low to accurately map the ROIs. | The overview has been taken in too low a magnification, or too small an area within a larger tissue section. | Take a brightfield image to capture large morphological details in the tissue or take a montage at higher magnification and stitch the images together. |
| 27 | There is no fluorescent signal for the antibody | a) The antibody penetration is poor.<br>b) the antibody penetration depth is limited<br>c) the antibody requires antigen retrieval.<br>d) The antibody does not bind the target of interest. | a) longer antibody incubation times are required.<br>b) antibody staining should be performed on thinner vibratome sections<br>c) Antigen retrieval may be required. We have tested limited heat-mediated antigen retrieval methods on human brain tissue without significant deterioration of the global ultrastructure (Supplementary Figure 9). However, this should be extensively tested for each case to assess the impact on the ultrastructural details of interest.<br>d) Test different antibodies for the same target. |
| 45 | Lead aspartate precipitates. | a) NaOH isn't fresh or the solution isn't at 60°C or pH is above 5.5.<br>b) Lead can precipitate with carbon dioxide. | a) Prepare fresh 1M NaOH solution and keep all solutions in a 60°C water bath during the pH adjustment and ensure the pH is adjusted only with NaOH and not corrected with acid if the pH goes above pH 5.5.<br>b) Use degassed solutions. |
| 57 | The image alignment is complicated due to tissue distortion or breakage during the embedding process. | The resin embedded sections are brittle or the resin embedding process causes shrinkage of the sections. | Only parts of the tissue that are closest to the ROIs need to be aligned precisely enough to draw the positions of the block to be excised. The BigWarp plugin in FIJI can be used to create the overlays with high precision (steps 116-125). |
| 63 | a) It is difficult to distinguish the blocks from each other on the tape after their excision from the section.<br>b) The section does not stick to the tape during excision of the laser cut blocks. | a) There are too many blocks with the same shape and/or they are in a different orientation to the empty regions left in the sections.<br>b) The ROI has not been cut completely. | a) Make distinctive shapes for the different blocks, cut identifying features within the block, and/or use binoculars to pick them up from tape.<br>b) Use tweezers to remove the section. |
| 88 | Bubbles have formed between the coverslip and the glass slide. | The air bubbles were formed when the coverslip was placed on the mounting medium | Before the glue dries, gently press the coverslip on the glass slide with the help of a pipette tip to remove the air bubbles. |
| 100 | There is a film over the sections that is not stained by the antibody, toluidine blue or hematoxylin. | The pathology pen was not completely dry before the antibody diluent was added to the slide causing a thin film of pen to form on top of the liquid surface. The film remains on the slide covering the sections when the liquid is tipped off preventing any further staining of the section | Ensure the pathology pen is completely air-dried before adding the antibody diluent to the slide. |
| 105 | The secondary antibody leaks all over the slide and isn't contained within the pathology pen. | The detergent from the PBS-T was not sufficiently removed from the surface of the slide. | Dry the surface of the pathology pen by rolling the end of a cotton bud over the pen boundary. Take care not to touch the sections. |
| 113 | No positive immunostaining was observed on the glass slides. | a) The staining conditions for the antibody need optimizing.<br>b) the distribution of the epitope is too diffuse to be observed in the ultra-thin sections | a) Test different antibody concentrations, incubation times and temperatures and common antigen retrieval methods.<br>b) cut thicker sections for the glass slides (e.g. 500 nm) to increase the number of epitopes available for staining in the sections. The etching time may also need to be increased accordingly. |
| 132 | Multiple features cannot be | The section was not | Ensure the free-floating |
|  | correlated between fluorescence and LM/EM. | completely flat during either the fluorescent imaging of the free-floating section, or the resin embedding causing large empty regions of empty resin within the ultra-thin sections impeding successful correlation, and/or the features that are visible on the individual ultrathin sections to be in different z-planes compared to what is anticipated based on the individual steps within the fluorescent z-stack. | section is completely flat under the coverslip before fluorescent imaging. Cut thicker vibratome sections to aid in the flat embedding. |
| 137 | Poor ultrastructure by EM. | The sample has not been properly fixed before resin embedding, or the sample has not been properly infiltrated during resin embedding | a) If using disease models, compare the ultrastructure to that of a control (non-diseased) sample prepared in the same way to ensure the ultrastructure is not disease related.<br>b) Post-fix the tissue with a higher amount of glutaraldehyde (up to 2.5%) after fluorescence imaging and before resin embedding.<br>c) Ensure there have been no errors in the resin-embedding process, that all steps were carefully followed, and the samples were not allowed to dry out between steps.<br>d) Increase infiltration times for the resin embedding protocol used. |
| 138 | Bad correlation between IHC/toluidine blue and EM. | The field-of-view of the EM micrograph is too limited for a good correlation. | We recommend taking two micrographs of each structure of interest: (1) an overview micrograph, i.e. a large FOV at low nominal magnification (typically around 630x) and (2) a detailed micrograph, i.e. a small FOV at high nominal magnification (typically |
|  |  |  | 8400x). Step 3 can then be split in two sub steps, namely. 3a. Toluidine blue (moving) to overview micrograph (target). 3b. Overview micrograph (moving) to detailed micrograph (target). |

#### Anticipated results

Using this protocol, it should be possible to perform CLEM on chemically fixed human brain tissue as well as other biological samples (Figure 9) resulting in the correlation of fluorescence, brightfield and EM images. The example below has been collected from the brain of a patient suffering from multiple systems atrophy with 4 hours post-mortem delay and fixed at autopsy in the fixation buffer for 75 hours. The fixed brain block was cut into 40 µm vibratome sections and immunostained with a neurofilament antibody cocktail to detect the neurons (neurofilament H, abcam ab5539; MAP2, abcam ab5392) and DAPI to detect cell nuclei (BioLegend cat # 422801). Following the creation of a fluorescent map, the sample was processed for electron microscopy, serial sectioning and electron microscopy image acquisition. The sections on glass slides were processed for IHC against alpha synuclein (pS129, abcam ab51253) and the subsequent alignment of fluorescence images, brightfield images (toluidine blue and IHC) and EM images enables us to correlate a single cell observed by fluorescence microscopy with its EM ultrastructure. Here, the correlation for a single neuron (n) has been done using four nearby nuclei (N) in the surrounding tissue. Combining a short post-mortem delay, optimized fixation solutions and correlation techniques enables us to achieve excellent preservation of ultrastructure (lipidic organelles) of the human brain tissue and optimal precision in defining ROIs. and from RT-CLEM and opening up new insights of the mechanisms behind the neurodegenerative diseases. This protocol should give consistent results in terms of ultrastructural quality for a given sample, if the sample was properly fixed and stored. Membranes, organelles, etc. should be well preserved in tissue using our inclusion criteria described previously.

**Figure 9.**
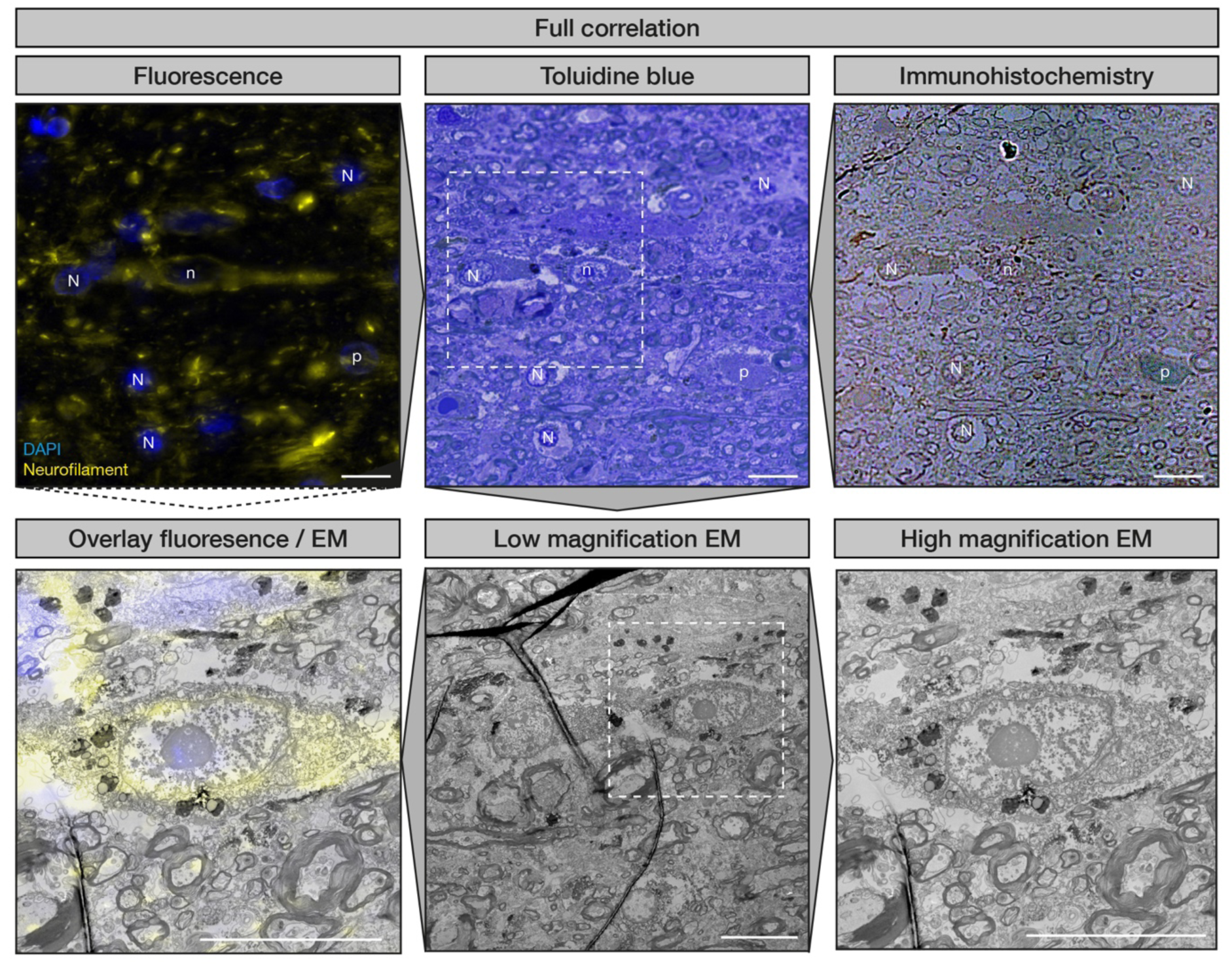
The full CLEM pipeline shown for the fluorescence and EM correlation of a single neuron within post-mortem human brain tissue. Fluorescent map generation and correlation to toluidine blue and IHC images in preparation for EM overlay. For correlation the same nuclei (N) are used that are visible in fluorescence, toluidine blue, and IHC. The target nucleus of the neuron desired for imaging is depicted with n. Typically, with IHC correlation immunostained regions of interest can be identified, as indicated by (p). (NB: not all nuclei observed in the fluorescence volume are visible in the single-plane LM images). Toluidine blue images are then correlated to low magnification EM images to identify the correct cells in EM. Based on the correlation of fluorescence with toluidine blue/IHC images and the correlation between toluidine blue/IHC images with EM images, fluorescence images can be overlaid with EM images with precision. These images were produced during the data collection from^11^. High magnification fluorescent z-stacks were deconvoluted using the THUNDER algorithm (Leica). Scale bar is 10 µm.

## Author contributions

AJL, SHS, CG, WvdB and HS conceived the strategy for carrying out CLEM on post-mortem human brain samples. WvdB collected, dissected and processed the brain tissue samples for CLEM. AJL and WvdB developed the strategy for fluorescent staining, imaging and mapping of free-floating brain sections with further optimization by DS, NS, and DB. AJL optimized the immunohistochemistry of resin sections on glass slides with contributions from MDF for the toluidine blue staining. CG optimized the manual *en bloc* staining and alternate serial sectioning protocol for 60 µm human brain sections. JD designed, printed and tested the sample holder components, and optimized the embedding protocol for the tissue auto-processor. LvdH wrote the macro to create .xml files. AJL and MDF optimized cutting of resin samples by laser-capture microdissection. DB and LvdH optimized the strategy for the image correlation of fluorescence and light microscopy with electron microscopy. CB and HS wrote the CLEM4CINA program. AJL tested the protocol on mouse brain sections and cell monolayers. NS, DS, MDF, DB, LvdH and AJL wrote the manuscript and prepared the figures. All authors approved the final version of the manuscript.

## Acknowledgments

We would like to thank the Netherlands brain bank autopsy team and Dr Jürgen Hench at the University Hospital Basel for the collection of post-mortem human brain samples used for the development of this protocol, Florent Pane from LGB,EPFL for printing the resin autoprocessor components, the Leica Research & Development team for assistance in the optimization of cutting resin samples with the Laser capture microdissection instrument, the staff at the Electron Microscopy Facility (UNIL) for training on equipment and TEMs and Dr. Ricardo Righetto (University of Basel) for assistance with the CLEM4CINA program. The final metal holders for the autoprocessor components were machined by Laser Automation, La-Chaux-de-Fond, Switzerland. AJL was supported by an EMBO short term fellowship (Grant no. 8878), the Synapsis Foundation Switzerland (Grant no. 2019-CDA01) and Parkinson Schweiz. This work was in part supported by the Swiss National Science Foundation (SNF Grants CRSII5_177195 and 310030_188548). WvdB is supported by the Dutch Parkinson association (Grant no. 2020-G01). Figure 1, Figure 3, Figure 6, and Supplementary Figure 7 were created with BioRender.com with agreement numbers CC275RDBFP, BF26TAAJ8K and GQ26TAAMU1 respectively.

## Data Availability

Original microscopy images in this manuscript can be found at BioImage Archive accession number S-BSST1770.

## Code Availability

All codes are available on github. CLEM4CINA.py can be accessed through the public repository with access under: https://github.com/LBEM-CH/clem4cina. The FIJI macro can be accessed through the public repository with access under: https://github.com/LBEM-CH/clem-for-human-brain.

## Supplementary Information

**Supplementary Table 1.** Antibodies compatible with 0.1% glutaraldehyde chemical fixation using immunofluorescence or IHC in human brain.

| Antibody target | Name (Clone) | Order information | FL dilution primary antibody | IHC |  |  | Reference of study tested in (FL; IHC) |
| --- | --- | --- | --- | --- | --- | --- | --- |
|  |  |  |  | Antigen retrieval conditions | Dilution primary antibody | Incubation time and temperature |  |
| alpha-Synuclein | LB509 | Thermofisher | 1/1000 | none | 1/500 | 1 hr / 37 °C | unpublished; 9-11, |
|  | aSyn (Clone 42) | BD Biosciences #15895639 | no signal | b | 1/100 | 4 hrs / RT | 10, 11, 13 |
|  | aSyn pS129 (EP1536Y) | Abcam #ab51253 | 1/5000 | none | 1/1000 | ON / 4 °C | 10 |
|  | aSyn pS129 (11a5) | Prothena | 1/1000 | none | 1/100,000 | 1 hr / 37 °C | 10; unpublished |
|  | aSyn C-Terminal Truncated x122 | Biolegend #A15127A | 1/1000 | a or b | 1/500 | ON / 4 °C | unpublished |
|  | aSyn aa 34-45 | Biolegend #15110D | 1/500 | a or b | 1/500 | ON / 4 °C | Unpublished |
| Astrocytes | GFAP | Abcam #4674 | 1/500 | n.d | n.d | n.d | 11 |
|  |  | Sigma-Aldrich #G3893 | 1/500 | n.d | n.d | n.d | unpublished |
|  |  | Agilent #Z033429-2 | 1/100 | n.d | n.d | n.d | 11 |
| Amyloid beta | beta-Amyloid (D54D2) XP(R) | Cell Signaling Technology #8243 | 1:200 | none | 1:1000 | 1 hr / 37 °C | unpublished |
| Lysosomes | LAMP1 | Abcam #ab24170 | 1/100 | n.d | n.d | n.d | unpublished |
|  | LIM11/Sr-B2 (SCARB2) | Novus Biologicals #NB400-129 | 1/800 | n.d | n.d | n.d | unpublished |
| Microglia | IBA1 | WAKO #019-19741 | 1/500 | n.d | n.d | n.d | 11 |
|  | CD45 (EP322Y) | Abcam #ab40763 | 1/100 | n.d | n.d | n.d | 11 |
|  | CD68 (KP1) * | BioRad # MCA1957 | 1/100 | n.d | n.d | n.d | 11 |
|  | P2RY12 | Sigma-Aldrich #HPA014518 | 1/100 | n.d | n.d | n.d | 11 |
|  | TMEM119 C-terminal | Abcam #ab185333 | 1/100 | n.d | n.d | n.d | 11 |
|  | TREM2 (D8I4C) | R&D Systems #MAB17291-100 | 1/100 | n.d | n.d | n.d | 11 |
| Mitochondria | TOMM20 (EPR15581-54) | Abcam #ab186735 | 1/100 | n.d | n.d | n.d | unpublished |
| Neurons | Neurofilament H | Sigma-Aldrich #ab5539 | 1/500 | n.d | n.d | n.d | 10, 11 |
|  | MAP2 | Abcam #ab5392 | 1/100 | n.d | n.d | n.d | 11 |
| Oligodendrocytes | MBP | LifeTechnologies #PA1-10008 | 1/100 | n.d | n.d | n.d | 11 |
|  | Olig2 | Sigma-Aldrich #ab9610 | 1/100 | n.d | n.d | n.d | 11 |
| p62 | SQSTM1 (cloneUMAB12) | VWR #ORIGUM500012 | no signal | none | 1/100 | 1 hr / 37 °C | 13 |
| phospho-Tau | pSer202, pThr205 (AT8) | Thermoshisher #MN1020 | 1:200 | none | 1:1000 | 4 hrs / RT | unpublished |
|  | pThr231 (AT180) | Thermoshisher #MN1040 | 1:500 | n.d. | n.d | n.d | unpublished |
|  | pS396 (E178) | Abcam #ab32057 | 1:500 | n.d. | n.d. | n.d. | unpublished |
|  | pS198 (EPR2400) | Abcam #ab79540 | 1:500 | n.d. | n.d. | n.d. | unpublished |
|  | pThr217 (44-744) | ThermoFisher #44-744 | 1:500 | n.d. | n.d. | n.d. | unpublished |
|  | pSer199 (2H23L4) | ThermoFisher #701054 | 1:500 | n.d. | n.d. | n.d. | unpublished |
|  | pSer396 (44-752G) | ThermoFisher #44-752G | 1:500 | n.d. | n.d. | n.d. | unpublished |
| Pre-cursor oligodendrocytes | NG2 | Abcam #ab255811 | 1/100 | n.d. | n.d. | n.d. | 11 |
| TDP43 | TDP43 (3H8) | Novus Biologicals | n.d. | none | 1/1000 | 1 hr / 37 °C | 12 |
| Tyrosine hydroxylase | Tyrosine hydroxylase | Sigma-Aldrich #AB152 | n.d. | none | 1/100 | ON/ 4 °C | unpublished |

FL= immunofluorescence; n.d.=no data; RT=room temperature; ON = overnight; FA = formic acid; hrs = hours; min = minutes; °C = degrees Celsius; \*stained processes only, a= 10 min in formic acid at RT and 30 min at 95 °C in sodium citrate pH 6, b= 10 min in formic acid at RT and 30 min at 95 °C in Tris-EDTA pH9.

**Supplementary Figure 1.**
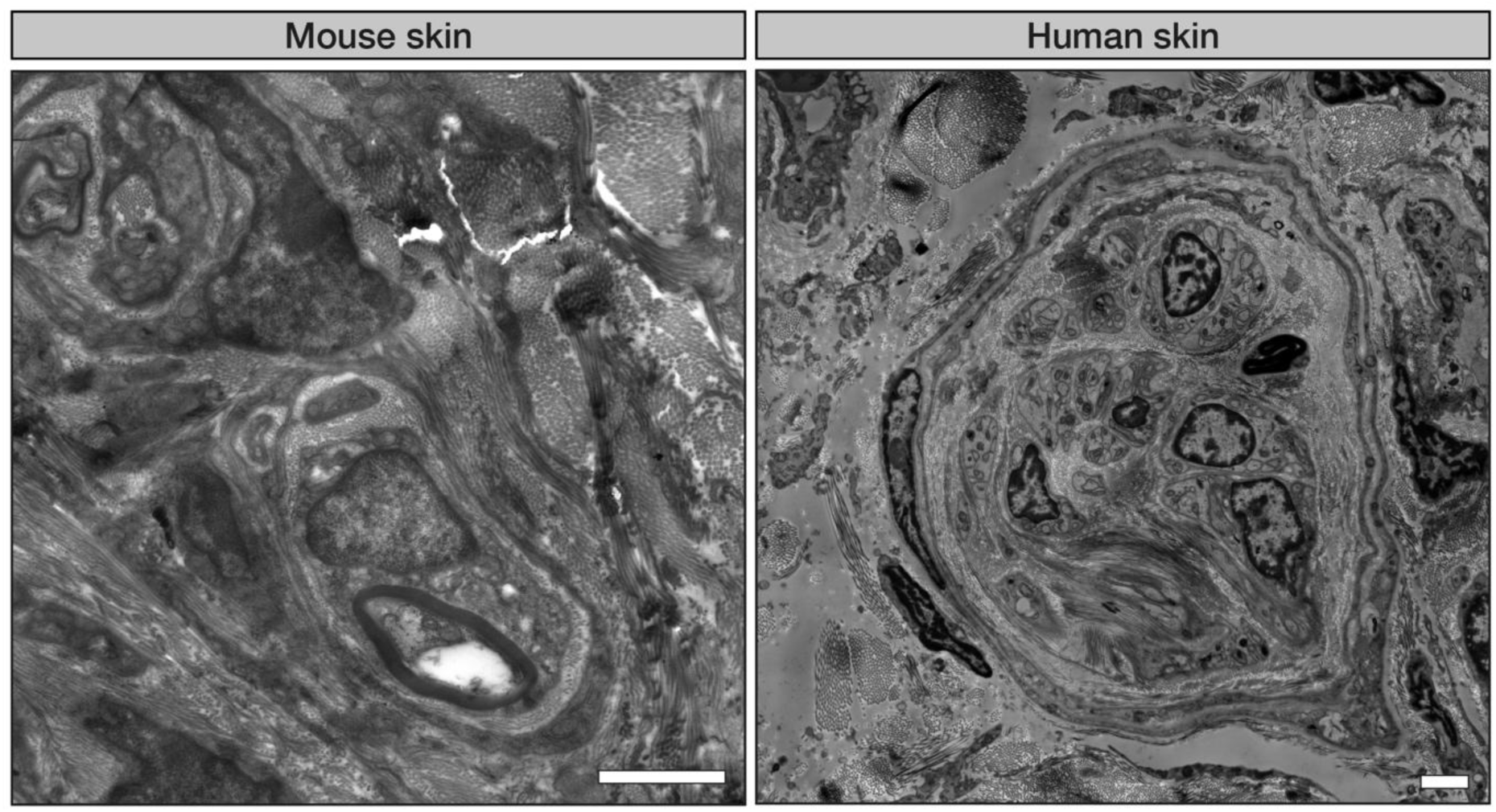
Example ultrastructure of mouse brain and human skin biopsy prepared using this pipeline. The ultrastructure of a dermal nerve fiber bundle from stained and resin embedded mouse skin biopsy. The mouse was perfused in 2% paraformaldehyde, 2.5% glutaraldehyde in 0.1 M phosphate buffer, pH 7.4. C - The ultrastructure of a dermal nerve fiber bundle from stained and resin embedded human skin tissue. 3 mm skin biopsy punches were collected at autopsy from the cervical C6-T1 region and fixed in 4% paraformaldehyde and 0.1% glutaraldehyde in 0.15 M cacodylate buffer. Both human and mouse skin biopsies were manually cut into 1 mm^3^ blocks and prepared for EM using the protocol for blocks in baskets. The images were taken at room-temperature on a Tecnai G2 Spirit (ThermoFisher) operated at 80 kV and equipped with a EMSIS Veleta camera. Scale bars are 2 µm.

### Protocol for autoprocessor program

Different resins can be used depending on the application and the sample. We routinely use Durcupan resin as it achieves the maximum hardness and is optimal for volume EM. However, when using the Leica tissue processor, EMBED 812 is preferred as the resin is less viscous and more efficiently infiltrates the tissue. We have provided the recipe we have used for both Durcupan and EMBED 812. Timing: 25 - 30 hours (depending on the program used, not including curing time)

## Materials

### Reagent

- Epon EMBED 812 embedding kit with BDMA (Electron Microscopy Sciences, cat. no. 14121) is supplied with 4 separate components: EMBED-812, DDSA, NMA and BDMA. CAUTION –Epon is toxic by inhalation, in contact with skin and if swallowed. Wear suitable clothing, gloves and eye protection.

### Reagent setup

#### EPON Resin without BDMA

- In glass beakers:

1. Prepare Mixture A: add 112.0 g EMBED-812 to 148.0 g DDSA.
2. Prepare Mixture B: add 103.0 g EMBED-812 to 94.0 g NMA.
3. 3. Add Mixture A to Mixture B and mix thoroughly by stirring slowly on a magnetic stirrer. Can be stored in syringes at −20°C.

EPON Resin with BDMA

- In glass beakers: take half (228.0 g) EPON Resin without BDMA previously prepared and add 1.5% w/w (3.4 g) BDMA and mix thoroughly without introducing air bubbles. Can be stored in syringes at −20°C.

Equipment

- Leica Tissue autoprocessor (Leica model EM TP)
- Camel hair paint brush (Electron Microscopy Sciences, cat. no. 65575-02)
- Custom sample carrier with sample disks
- Standard sample baskets for large samples/tissue blocks
- Plastic autoprocessor tubes
- Syringes

Procedure

The programs for free-floating sections and large blocks are provided. The full protocol for free-floating sections is split into two programs with the lead aspartate staining done manually at 60°C.

1. Add the correct number of empty tubes to the rotation device depending on the protocol to be used.
2. Put the empty sample carrier in the autoprocessor.
3. Check that the processor can rotate nicely into every tube without catching.
4. Place an empty sample disk (flat side down) into the sample holder and immerse into the buffer.
5. Place a section on top of the sample disk and cover with another disk (side down).
6. Continue placing sections and stacking disks into the carrier as needed.
7. Add the correct spacer and assemble the rest of the carrier.
8. Leave the sample carrier immersed in the buffer.
9. Prepare the solutions required and add them to the tubes leaving the alcohol until last to avoid evaporation.
10. Insert the carrier into the autoprocessor
11. Start the program.

NOTE-Here, we use acetone rather than ethanol for dehydration and embedding but both can be used depending on the tissue and resin used. Our effort has been to avoid using propylene oxide, therefore using acetone preferentially.

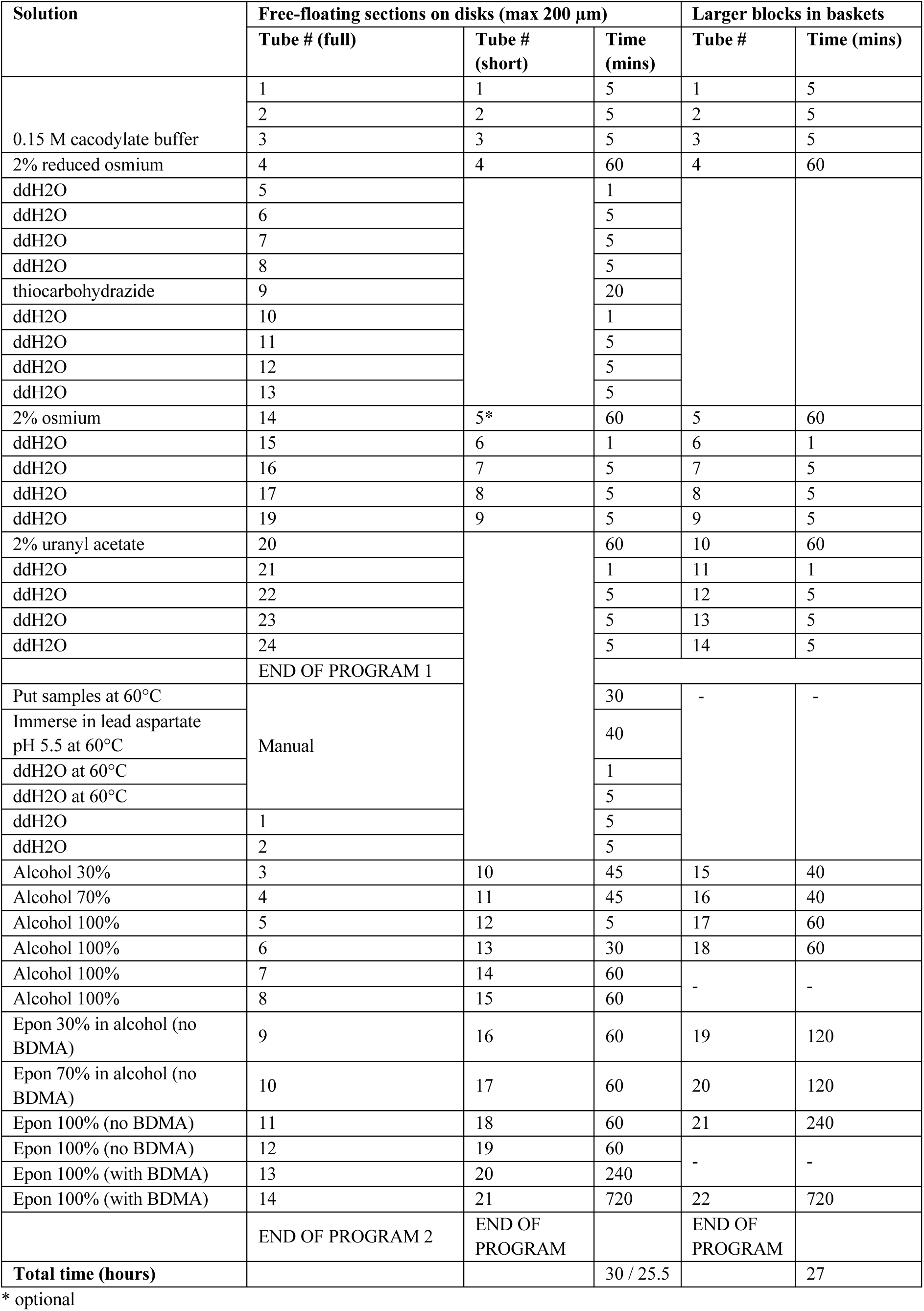

Autoprocessor sample holder

**Supplementary Figure 2.**
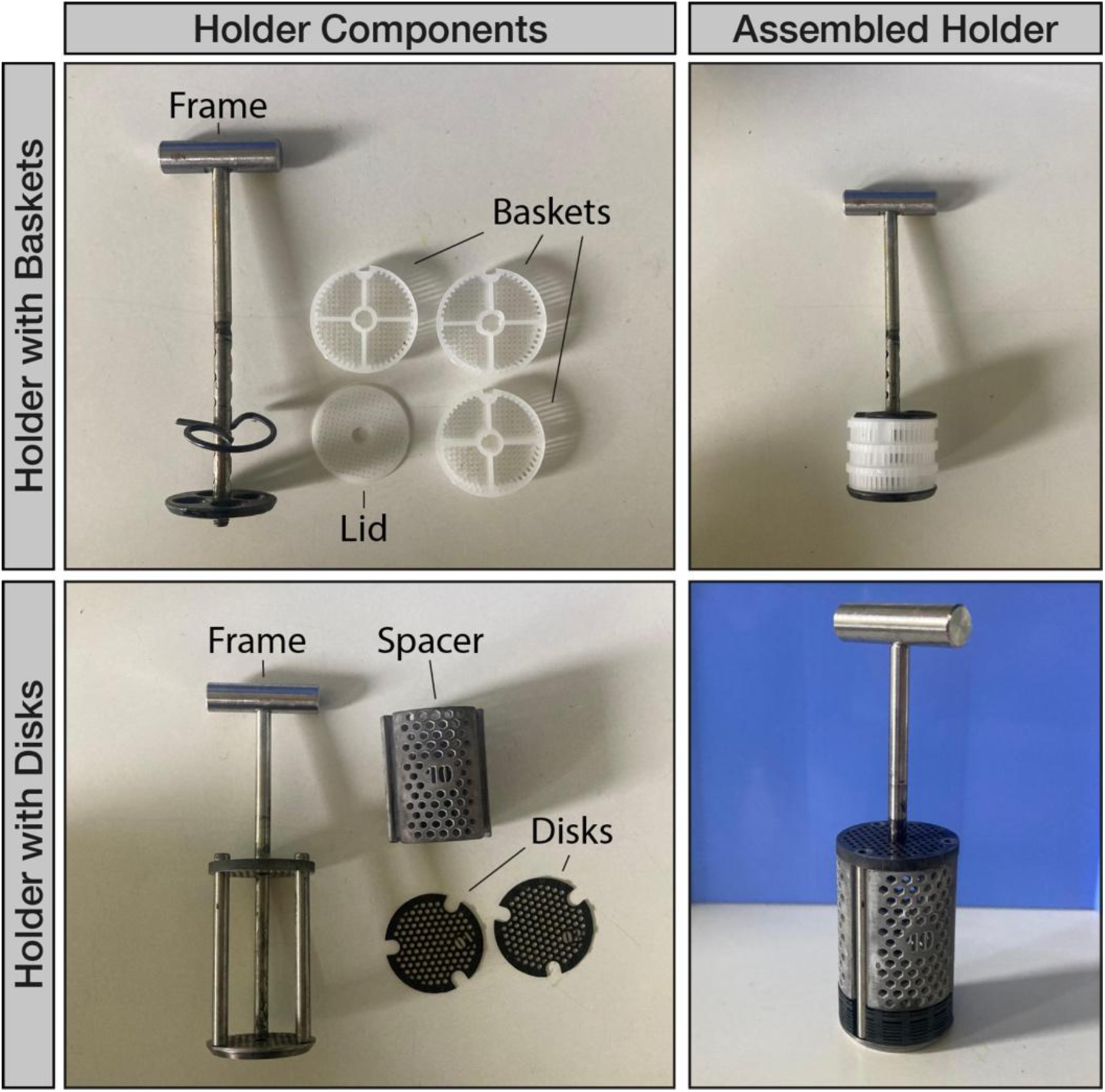
Autoprocessor sample holder for resin embedding of thin sections.

The samples are processed in either the provided sample holder with baskets for thick sections (top) or in custom-made metal disks for free floating sections (bottom).

Schematic for custom autoprocessor sample holder

**Supplementary Figure 3.**
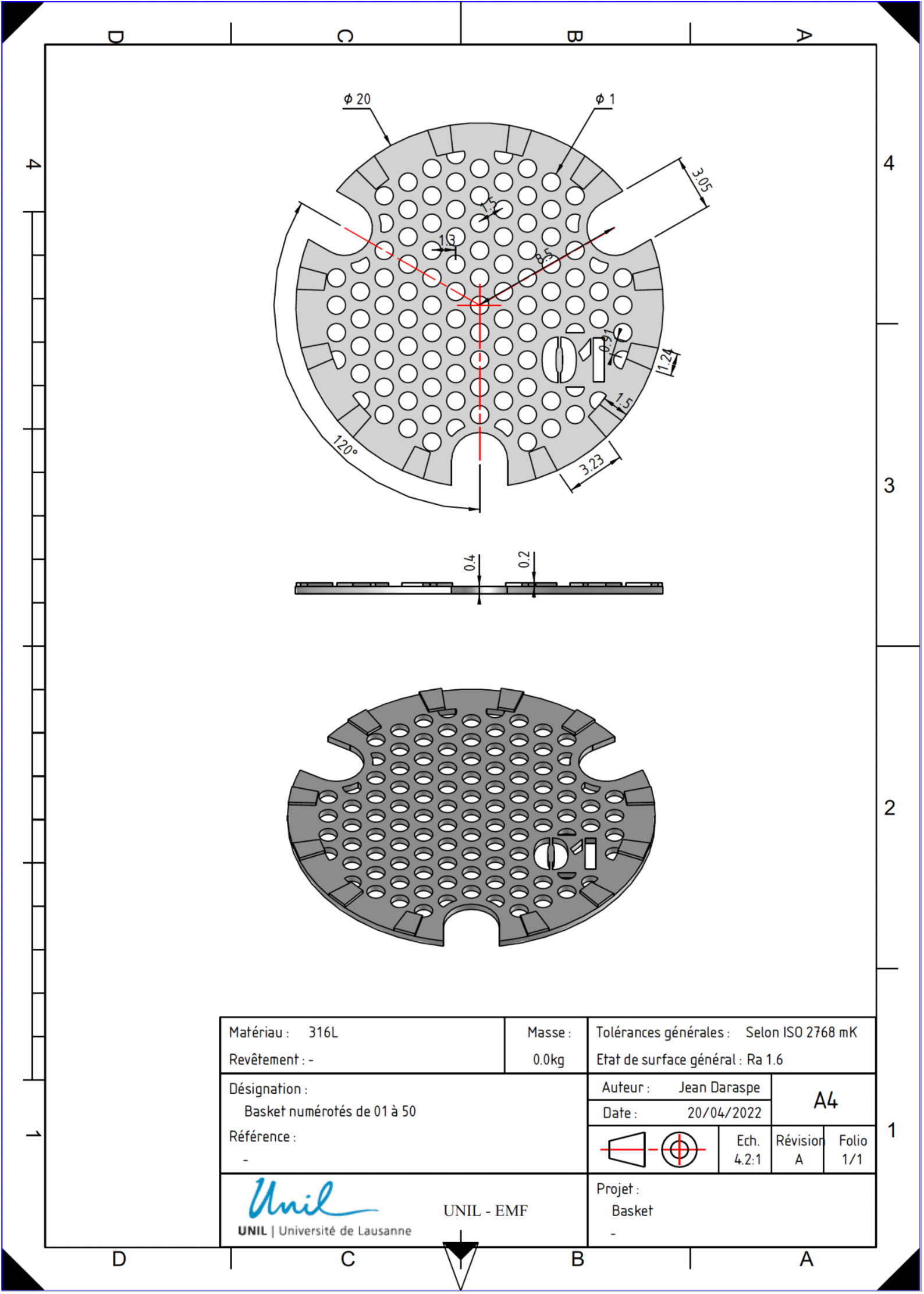
Schematics for the printing of custom-made sample disks for use with a tissue autoprocessor. These schematics can be used for 3D printing with an alcohol-resistant resin however we recommend having them manufactured in surgical steel. Units in mm.

**Supplementary Figure 4.**
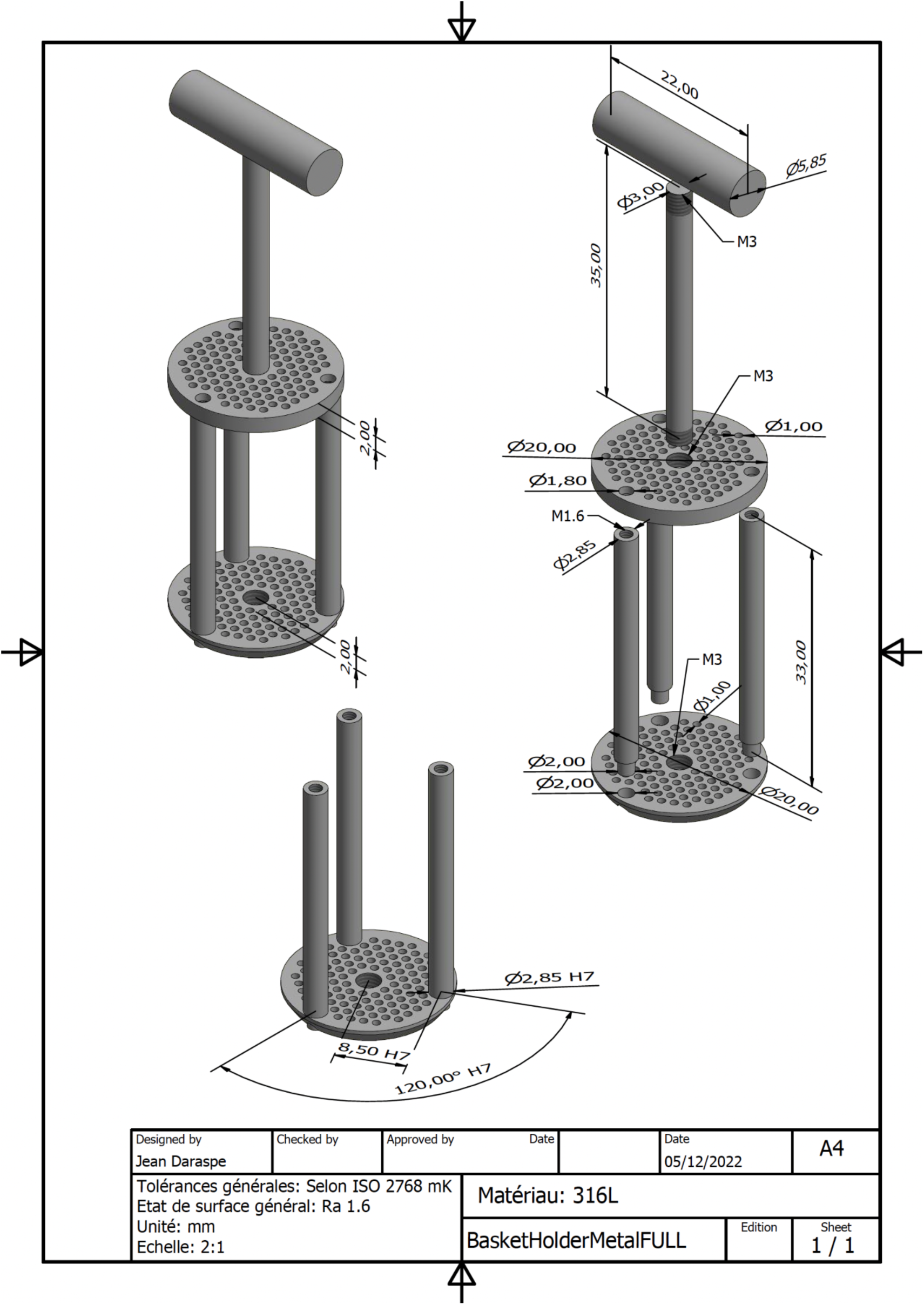
Schematics for printing the sample holder to house the custom-made sample holder. These schematics can be used for 3D printing with an alcohol-resistant resin however we recommend having them manufactured in surgical steel. Units in mm.

Laser microdissection

Timing: 1 hour (depending on number of ROI’s to be cut)

Materials and Equipment

- Leica Laser microdissection (LMD7)
- Double-sided tape
- OPTIONAL LMD_macro_for_FIJI (https://github.com/LBEM-CH/clem-for-human-brain)

Procedure

An xml file containing the XY coordinates for user defined shapes can be created manually (A) or using an automation script (B).

1. Peel the embedded sections off the Aclar sheet and mount on a glass slide using double sided tape (Supplementary Figure 5A). NOTE-do this carefully as the resin can easily tear through the embedded section.
2. Load glass slide into the slide holder of a brightfield microscope. Make 3 calibration crosses with laser into the resin surrounding the section (top left, top right, bottom right) using the 4X objective.
3. Take a brightfield image of the section at a magnification suitable to visualize the calibration marks.
4. Overlay this image with the fluorescent map and annotate the areas to be laser cut.
5. Export the image of the section with the drawn shapes and open it in FIJI.
6. Prepare the .xml file manually (A) or using our custom CLEM-for-human-brain macro (B)

A. Manually (Supplementary Figure 5B):

- i. use the multipoint tool to select the center of each calibration point and the corners of each shape with as much precision as possible.
- ii. Click > Analyze > Measure (or Cmd M) to obtain the xy coordinates of each point clicked
- iii. Enter these into a .txt file using the format supplied (Supplementary Fig 5) CRITICAL: use whole numbers only. Round where necessary
- iv. Change the extension of the .txt file to .xml.

B. LMD macro (Supplementary Figure 5C)

- i. Download the CLEM-for-human-brain macro (see Material) CRITICAL: macro requires FIJI version 1.53d or later.
- ii. Open the macro in FIJI by dragging and dropping it into the FIJI bar
- iii. Click ‘run’.
- iv. Follow the series of pop-ups asking you to set some parameters, mark the calibration points, and mark the corners of the shapes.
- v. A .xml file (Supplementary Figure 6) will be saved in the directory indicated at the start.

1. Import the .xml file into the microscope software.
2. Follow the prompts for sample calibration.
3. Check that all shapes have been imported and are in the correct position.
4. Start laser cutting

**Supplementary Figure 5.**
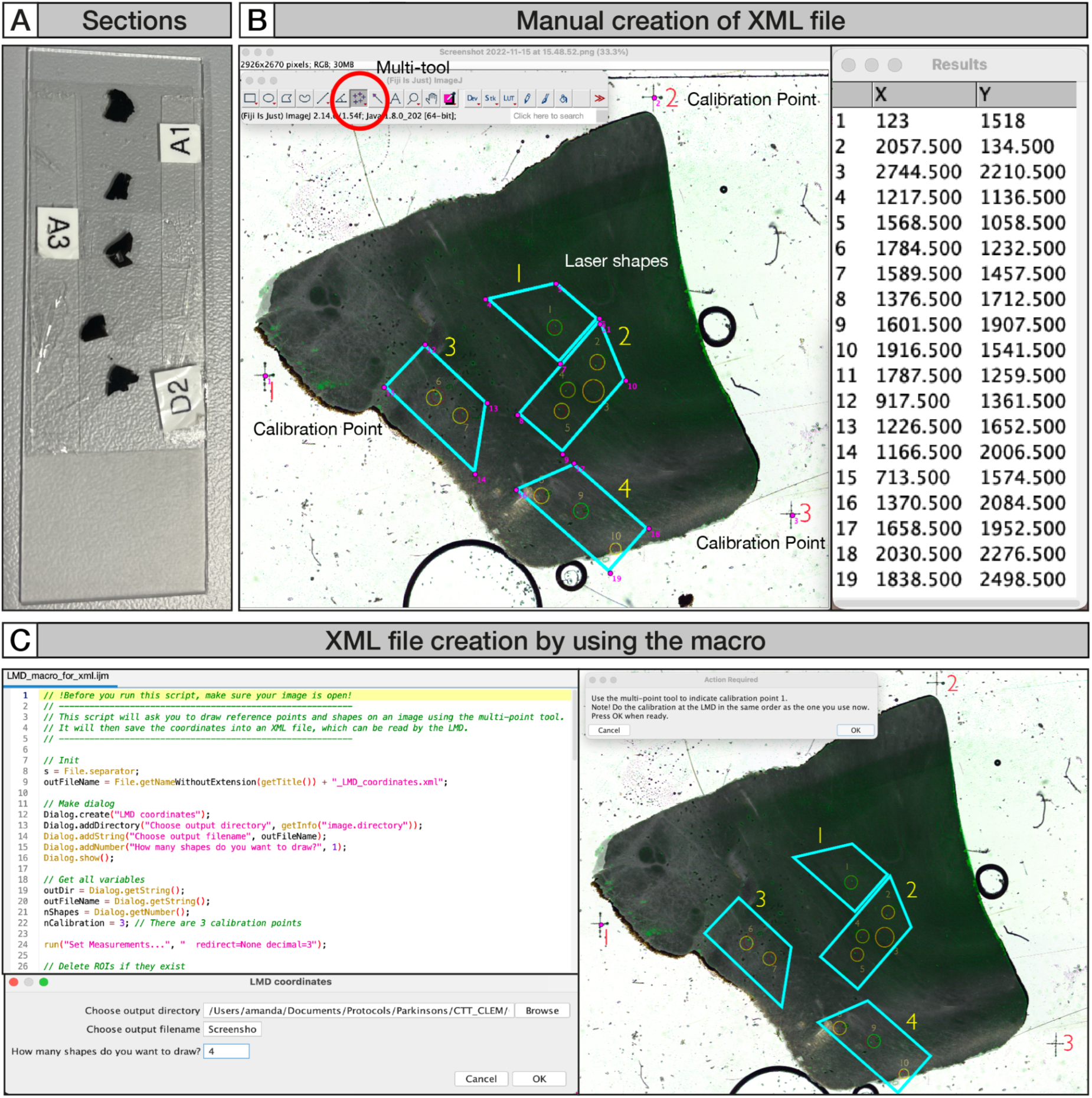
Excision of ROIs by laser cutting with the Leica LMD. A) The resin embedded sections are mounted on glass slides between two strips of double-sided tape; B) Steps for the manual creation of an xml file. The multi-point tool is used to select the center of each calibration point and the corners of each shape. The resulting XY coordinates are manually entered into a txt file; C) creation of the xml file with the LMD macro. The macro is opened and run in FIJI. Pop-ups guide the user through the addition of points using the multi-point tool. An xml file is automatically created.

**Supplementary Figure 6.**
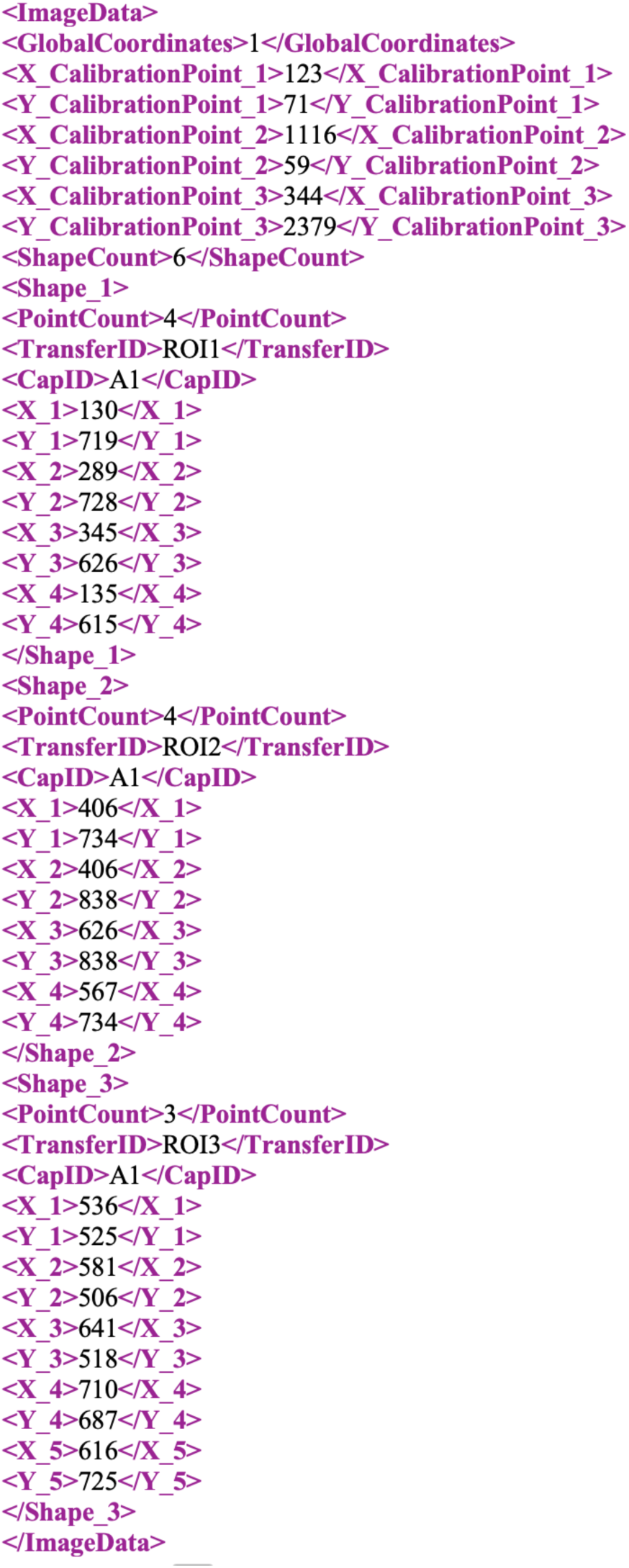
Example of the text and format of the xml file required for the import of shapes to the Leica LMD7. All information between the >< symbols (e.g. The XY coordinates, number of shapes and point count) can be entered manually or created automatically with the CLEM-for-human-brain macro.

Hematoxylin counterstaining

Timing: 1.5 hours

Materials

Reagents

- Hematoxylin (Mayer) for histology (J.T. Baker, cat. no. 3870.1000)
- CAUTION-Hematoxylin can cause irritation to the eyes and skin and it is toxic if swallowed. Wear suitable clothing, gloves and eye protection.
- Clearing solution, NeoClear (Sigma-Aldrich, cat. no. 1.09843.5000)

Reagent setup

1% HCl and 70% Ethanol in ddH2O (Acid-Ethanol)

- Mix 5.0 ml of HCl with 0.35 L Ethanol make up the final volume to 0.5 L with ddH2O. CAUTION-Work with HCl in a fume hood at all times.

Procedure

1. Place all the slides on a glass slide staining rack and fill all the wells according to this order:

1: Hematoxylin

2: ddH2O

3: Tap water

4: Acid-Ethanol

5 and 6: ddH2O

7: Tap water

8-10: 100% Ethanol

11-12: Clearing solution

1. Counterstain slides in well (1) for 5-10 min. NOTE-The timing should be optimized depending on the sample used. We have found that most tissues stain within 5 mins. Stain cell monolayers for 10 mins.
2. Rinse slides in well (2) for a few seconds.
3. Wash slides in well (3) for 5 mins.
4. Rinse slides in well (4) for a few seconds.
5. Wash slides in wells (5) and (6) for 1 min each.
6. Wash slides in well (7) for 2 min.
7. Rinse slides in wells (8), (9) and (10) for a few seconds.
8. Rinse slides in wells (11) and (12) for a few seconds.

**Supplementary Figure 7.**
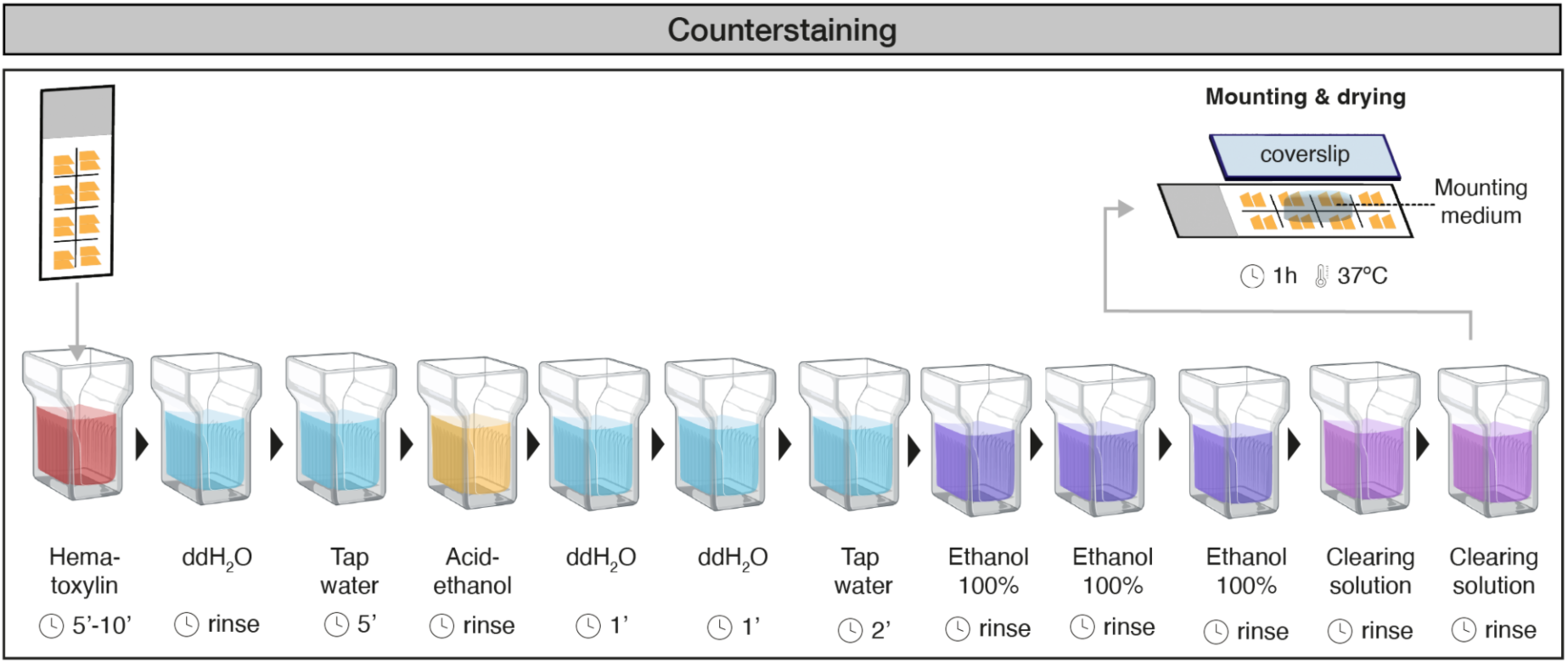
Schematic for hematoxylin staining.

CLEM4CINA

The code can be download from: https://github.com/LBEM-CH/clem4cina

Procedure

The generation of the correlation matrix works by filling in three X,Y coordinates into the corresponding entry widgets (red frame) and pressing the ‘Prepare LM2EM’ and/or ‘Prepare EM2LM’ button widgets. Any LM or EM X,Y coordinate can then be converted into the other respective coordinate system by filling in the entry widgets in the green frame and pressing the ‘LM->EM’ or ‘LM<-EM’ button widgets (Supplementary Figure 8).

1. Download and install the program
2. Run CLEM4CINA from the applications folder

- a. Python clem4cina.py
3. Click on Load image
4. Click the “Control points” button on the pop-up window
5. Click on three control points in the LM image (e.g., the corners of the section)
6. Drive to the same control points on the TEM
7. Enter the XY stage coordinates in the entry widgets for Control Points > EM
8. Click Prepare LM2EM
9. Click the “5 ROIs” button on the LM image
10. Click on up to 5 ROIs in the LM image
11. Click LM->EM in the GUI
12. Enter the resulting XY coordinates into the TEM software to drive to that stage position.

**Supplementary Figure 8.**
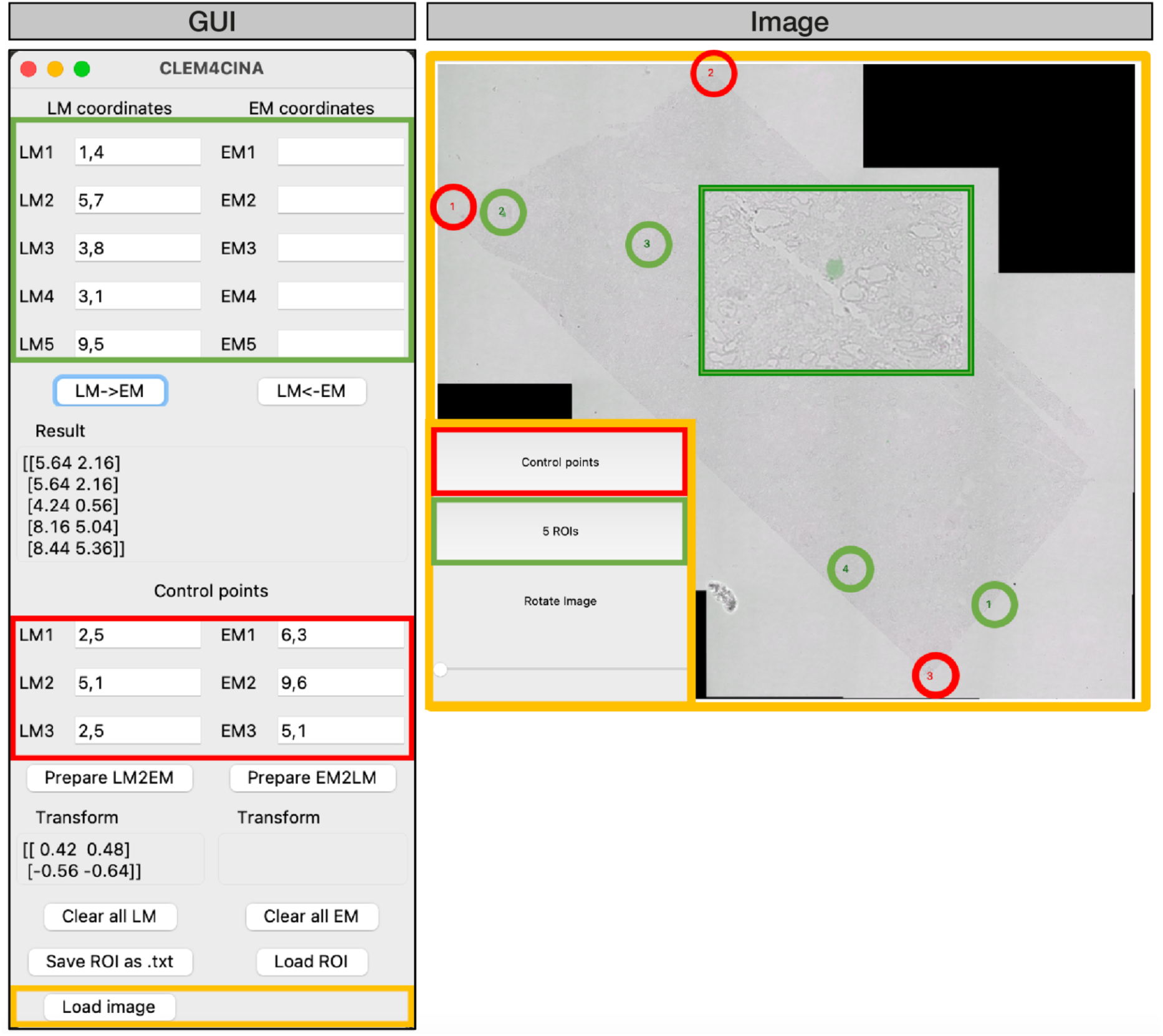
The GUI for the CLEM4CINA program. An image is loaded into the GUI (orange frame GUI) that will then be displayed in a top view window (orange frame, Image). Two button widgets (inset, Image) prompt the user to click on three control points (red frame) and five ROIs (green frame) in the loaded image. Examples of selected points are displayed and numbered in the image in the corresponding colours. The coordinates are automatically entered into the green and red frames of the main GUI. The mouse wheel can be used to zoom into the image (green frame, left). A slider on the widget panel under ‘Rotate Image’ (inset, Image) allows the user to rotate the displayed image.

**Supplementary Figure 9.**
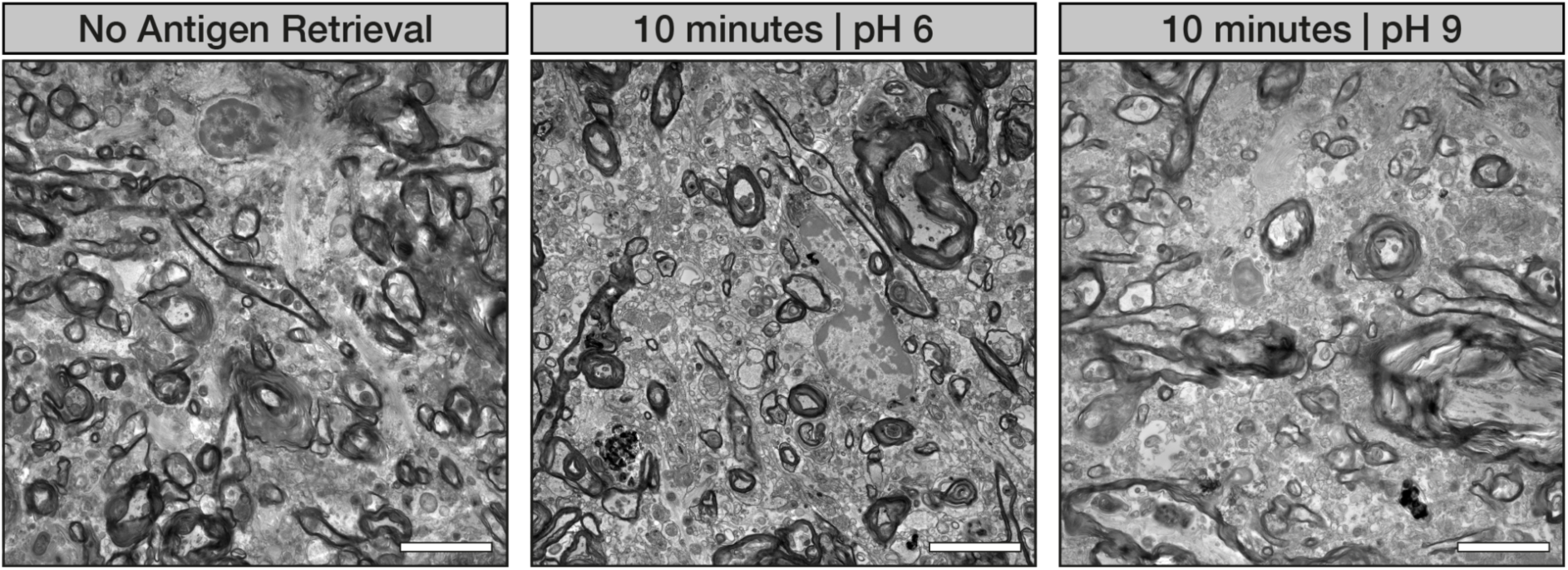
Comparison of human brain tissue ultrastructure for three different antigen retrieval conditions. No alteration in the tissue ultrastructure was visible in all the conditions. No antigen retrieval, heat-mediated antigen retrieval for 10 min with citrate buffer (pH 6) at 95°C, and heat-mediated antigen retrieval for 10 min with tris-EDTA buffer (pH 9) at 95°C. TEM images were taken with a Tecnai Spirit BioTwin (ThermoFisher) operated at 80 kV and equipped with a TVIPS F416 camera. Scale bar is 5 µm.

## References

1. Lewis, A. J. et al. Imaging of post-mortem human brain tissue using electron and X-ray microscopy. Current Opinion in Structural Biology 58, 138–148 (2019).

2. Genoud, C., Titze, B., Graff-Meyer, A. & Friedrich, R. W. Fast Homogeneous En Bloc Staining of Large Tissue Samples for Volume Electron Microscopy. Front. Neuroanat. 12, (2018).

3. Hua, Y., Laserstein, P. & Helmstaedter, M. Large-volume en-bloc staining for electron microscopy-based connectomics. Nat Commun 6, 7923 (2015).

4. Tapia, J. C. et al. High-contrast en bloc staining of neuronal tissue for field emission scanning electron microscopy. Nat Protoc 7, 193–206 (2012).

5. Deerinck, T. J., Bushong, E. A., Ellisman, M. H. & Thor, A. Preparation of Biological Tissues for Serial Block Face Scanning Electron Microscopy (SBEM). (2022).

6. Tanner, H., Sherwin, O. & Verkade, P. Labelling strategies for correlative light electron microscopy. Microscopy Research and Technique 86, 901–910 (2023).

7. de Boer, P., Hoogenboom, J. P. & Giepmans, B. N. G. Correlated light and electron microscopy: ultrastructure lights up! Nat Methods 12, 503–513 (2015).

8. Müller, A. et al. Modular segmentation, spatial analysis and visualization of volume electron microscopy datasets. Nat Protoc 19, 1436–1466 (2024).

9. Shahmoradian, S. H. et al. Lewy pathology in Parkinson’s disease consists of crowded organelles and lipid membranes. Nature Neuroscience 22, 1099–1109 (2019).

10. Lewis, A. J. et al. Ultrastructural diversity of alpha-Synuclein pathology in the post-mortem brain of Parkinson patients: implications for Lewy Body formation. *bioRxiv* 2024.07.25.605088 (2024) doi:10.1101/2024.07.25.605088.

11. Böing, C. et al. Distinct ultrastructural phenotypes of glial and neuronal alpha-synuclein inclusions in multiple system atrophy. Brain awae137 (2024) doi:10.1093/brain/awae137.

12. De Rossi, P. et al. FTLD-TDP assemblies seed neoaggregates with subtype-specific features via a prion-like cascade. EMBO reports 22, e53877 (2021).

13. Lam, I. et al. Rapid iPSC inclusionopathy models shed light on formation, consequence, and molecular subtype of α-synuclein inclusions. Neuron (2024) doi:10.1016/j.neuron.2024.06.002.

14. Burger, D. et al. 1.94 Å structure of synthetic α-synuclein fibrils seeding MSA neuropathology. 2024.07.01.601498 Preprint at 10.1101/2024.07.01.601498 (2024).

15. Karreman, M. A. et al. Fast and precise targeting of single tumor cells in vivo by multimodal correlative microscopy. J Cell Sci 129, 444–456 (2016).

16. Collinson, L. M., Carroll, E. C. & Hoogenboom, J. P. Correlating 3D light to 3D electron microscopy for systems biology. Current Opinion in Biomedical Engineering 3, 49–55 (2017).

17. Lane, R., Wolters, A. H. G., Giepmans, B. N. G. & Hoogenboom, J. P. Integrated Array Tomography for 3D Correlative Light and Electron Microscopy. Front Mol Biosci 8, 822232 (2022).

18. Kolotuev, I. Work smart, not hard: How array tomography can help increase the ultrastructure data output. J Microsc (2023) doi:10.1111/jmi.13217.

19. Micheva, K. D. & Smith, S. J. Array tomography. Neuron 55, 25–36 (2007).

20. Kopek, B. G. et al. Diverse protocols for correlative super-resolution fluorescence imaging and electron microscopy of chemically fixed samples. Nat Protoc 12, 916–946 (2017).

21. Maco, B. et al. Semiautomated correlative 3D electron microscopy of in vivo-imaged axons and dendrites. Nat Protoc 9, 1354–1366 (2014).

22. Liu, M. et al. Tracking endocytosis and intracellular distribution of spherical nucleic acids with correlative single-cell imaging. Nat Protoc 16, 383–404 (2021).

23. Darcy, K. J., Staras, K., Collinson, L. M. & Goda, Y. An ultrastructural readout of fluorescence recovery after photobleaching using correlative light and electron microscopy. Nat Protoc 1, 988–994 (2006).

24. Sabatini, D. D., Bensch, K. & Barrnett, R. J. Cytochemistry and electron microscopy. The preservation of cellular ultrastructure and enzymatic activity by aldehyde fixation. J Cell Biol 17, 19–58 (1963).

25. Morin, F., Crevier, C., Bouvier, G., Lacaille, J.-C. & Beaulieu, C. A Fixation Procedure for Ultrastructural Investigation of Synaptic Connections in Resected Human Cortex. Brain Research Bulletin 44, 205–210 (1997).

26. Schultz, R. L. & Willey, T. J. Extracellular space and membrane changes in brain owing to different alkali metal buffers. J Neurocytol 2, 289–303 (1973).

27. August, B. K., Kong, L., Massey, R. J., Zhang, S.-C. & Strader, T. E. Robotic Optimization of Specimen Preparation Protocol for Astrocytes Seeded on Coverslips for Imaging by Transmission Electron Microscopy (TEM). Microscopy and Microanalysis 29, 1106–1110 (2023).

28. Kuipers, J. & Giepmans, B. N. G. Neodymium as an alternative contrast for uranium in electron microscopy. Histochem Cell Biol 153, 271–277 (2020).

29. Bogovic, J. A., Hanslovsky, P., Wong, A. & Saalfeld, S. Robust registration of calcium images by learned contrast synthesis. in 2016 IEEE 13th International Symposium on Biomedical Imaging (ISBI) 1123–1126 (2016). doi:10.1109/ISBI.2016.7493463.

30. Schindelin, J., et al. Fiji: an open-source platform for biological-image analysis. *Nat Methods* 9, 676–682 (2012).

31. Paul-Gilloteaux, P. et al. eC-CLEM: flexible multidimensional registration software for correlative microscopies. Nat Methods 14, 102–103 (2017).

32. Navarro, P. P. et al. Cerebral Corpora amylacea are dense membranous labyrinths containing structurally preserved cell organelles. Scientific Reports 8, 18046 (2018).

33. Tamada, H., Blanc, J., Korogod, N., Petersen, C. C. & Knott, G. W. Ultrastructural comparison of dendritic spine morphology preserved with cryo and chemical fixation. Elife 9, e56384 (2020).

34. Korogod, N., Petersen, C. C. & Knott, G. W. Ultrastructural analysis of adult mouse neocortex comparing aldehyde perfusion with cryo fixation. eLife 4, e05793 (2015).

35. Griffiths, G. Fine Structure Immunocytochemistry. (Springer Science & Business Media, 2012).

36. Mollenhauer, H. H. Artifacts caused by dehydration and epoxy embedding in transmission electron microscopy. Microscopy Research and Technique 26, 496–512 (1993).

37. Schnell, U., Dijk, F., Sjollema, K. A. & Giepmans, B. N. G. Immunolabeling artifacts and the need for live-cell imaging. Nat Methods 9, 152–158 (2012).

38. Yi, H., Leunissen, J., Shi, G., Gutekunst, C. & Hersch, S. A novel procedure for pre-embedding double immunogold-silver labeling at the ultrastructural level. J Histochem Cytochem 49, 279–284 (2001).

39. Morikawa, S., Sato, A. & Ezaki, T. A simple, one-step polychromatic staining method for epoxy-embedded semithin tissue sections. Microscopy (Oxf) 67, 331–344 (2018).

